# Element content and distribution has limited, tolerance metric dependent, impact on salinity tolerance in cultivated sunflower (*Helianthus annuus*)

**DOI:** 10.1101/2019.12.11.872929

**Authors:** A.A. Temme, V.A. Burns, L.A. Donovan

## Abstract

Disruption of ion homeostasis is a major component of salinity stress’s effect on crop yield. In cultivated sunflower prior work revealed a trade-off between vigor and salinity tolerance. Here we determined the association of elemental content/distribution traits with salinity tolerance, both with and without taking this trade-off into account. We grew seedlings of twelve *Helianthus annuus* genotypes in two treatments (0/100 mM NaCl). Plants were measured for biomass (+allocation), and element content (Na, P, K, Ca, Mg, S, Fe, B, Mn, Cu, Zn) in leaves (young and mature), stem, and roots. Genotype tolerance was determined by the proportional decline in biomass and as the deviation from the expected vigor/tolerance trade-off. Genotype rankings on these metrics were not the same. Elemental content and allocation/distribution were highly correlated both at the plant and organ level. Suggestive associations between tolerance and elemental traits were fewer and weaker than expected and differed by tolerance metric. Given the highly correlated nature of elemental content, it remains difficult to pinpoint specific traits underpinning tolerance. Results do show that taking vigor related trade-offs into account is important in determining traits related to tolerance and that the multivariate nature of associated traits should be considered.

## Introduction

High soil salinity is a major abiotic stress impacting crop yield worldwide. Due to natural processes or as the result of land clearing, unsustainable irrigation practices, and high evaporation high levels of salt (generally NaCl) are present in the soil (Munns and Gilliham, 2015). In these salinized environments, plants experience the twin stresses of a high soil osmotic potential, limiting water uptake, and a high concentration of sodium, leading to ionic stress (Munns, 2002). This ionic stress is caused by plants taking up sodium at the expense of essential elements such as potassium and calcium (Hawkesford *et al.*, 2012). The accumulation of Na to toxic levels, in combination with elemental imbalances within tissues, disrupts cellular and enzymatic functions (Chapin, 1980; Negrão *et al.*, 2017; Morton *et al.*, 2018). With 20% of the worlds irrigated agricultural land affected by salinity (FAO 2005), it is imperative that we improve crop salt tolerance to continue feeding a growing global population.

The effect of salt stress on plant elemental status has been a central focus of research. Given sodium’s central role in salt stress, lower Na concentrations in shoot tissues has been shown to correlate with salt tolerance in crops (Flowers and Yeo, 1995; Flowers, 2004). Additionally, given the antagonistic effect of high sodium concentrations in the soil on potassium uptake, maintenance of potassium levels is correlated with tolerance as well (Akram *et al.*, 2009; Hawkesford *et al.*, 2012; Demidchik, 2014; Temme *et al.*, 2019). However, most of these studies are limited to leaf or shoot versus root comparisons, restricting analyses to correlations between individual elemental concentrations and salt tolerance. By taking a whole-plant and multiple tissue approach, and incorporating a broader range of elements, we can infer active mechanisms of elemental allocation, such as exclusion or accumulation (Abrahamson and Caswell, 1982; Romero and Maranon, 1996) and relate those mechanism to salt tolerance.

While the role of sodium and potassium during salinity stress has been extensively studied (Shabala and Cuin, 2008; Shabala, 2017; Wu *et al.*, 2018) less focus has been placed on other elements (Broadley *et al.*, 2012; Lewis, 2018). Over the past decade inductively coupled plasma mass spectroscopy (ICP-MS) has opened up the door for relatively inexpensive analysis of elemental content of plant tissue (Salt *et al.*, 2008), leading to the identification of a host of genes involved in plant elemental content (Whitt *et al.*, 2018). In previous work on cultivated sunflower ICP-MS analysis revealed leaf sulfur content, besides potassium, to play a role in salinity tolerance (Temme *et al.*, 2019). Given the impacts of salinity stress on plant nutrient stoichiometry, a comprehensive picture of plant elemental content has the potential to shed light on mechanisms of salt sensitivity and tolerance (Negrão *et al.*, 2017).

Stress tolerance can be difficult to quantify and numerous metrics for tolerance exist (Zhu *et al.*, 2016; Morton *et al.*, 2018)). Generally, salinity tolerance is defined as having a high relative performance (i.e. low proportional decline in performance between control and saline conditions, hereafter “proportional-decline-tolerance” metric) (Munns and Tester, 2008; Negrão *et al.*, 2017). An ideal crop cultivar would combine this low proportional decline with high vigor under benign conditions. However, studies have shown there can be a tradeoff between growth and stress tolerance (Mayrose *et al.*, 2011; Koziol *et al.*, 2012), and indeed for cultivated sunflower there appears to be a negative trade-off between vigor (growth in benign conditions) and the proportional effect of salt stress. More vigorous genotypes have a greater proportional decline in biomass under salt stress (Temme *et al.*, 2019). Given this trade-off, selecting for the ideal cultivar with high vigor and high tolerance is challenging using the proportional-decline-tolerance metric.

Fortunately, it is possible to construct an additional tolerance metric that takes this vigor/”tolerance” trade-off into account. By scoring genotypes on their deviation from the expected effect of salinity stress based on their biomass, we can redefine tolerance by taking vigor into account. This metric allows us to disentangle traits related to vigor from traits related to tolerance (performing better than expected under stress). Finding traits related to this metric of tolerance could allow us to incorporate them in vigorous genotypes to select for the ideal high-vigor/high-tolerance genotype. However, the question remains on whether this “expectation-deviation-tolerance” relates to the more conventional proportional-decline-tolerance metric.

Cultivated sunflower (*Helianthus annuus*), a major oilseed crop, is a moderately salt tolerant species (Katerji et al. 2000), and previous studies have shown there is genotypic variation in response to salt stress and elemental status (Ashraf and Tufail, 1995; Shi and Sheng, 2005; Temme *et al.*, 2019). We ask the following questions in order to determine the effect of taking a known vigor/salinity effect trade-off into account when determining tolerance and develop a more comprehensive picture of the effects of salt stress on elemental status and tolerance. (1) Do genotypes differ in the effect of salinity on biomass and how do genotypes compare for salinity tolerance measured as either proportional-reduction-tolerance or expectation-deviation-tolerance? (2) Are there genotypic differences in the effect of salinity on whole plant and organ (leaf, stem, root) level elemental content? (3) Are whole plant and organ level elemental content and allocation associated with either metric of salinity tolerance?

## Materials and Methods

### Plant Growth

Achenes from twelve genetically diverse lines of *H. annuus* (Table 3.S1) were germinated at the University of Georgia greenhouses in a 3:1 sand to turface MVP® (Turface Athletics, PROFILE Products, LLC, Buffalo Grove, IL) mixture. Ten days after germination, seedlings were transplanted into 30 cm tall, 5L pots filled with the same growth substrate, supplemented with 15ml of lime (Austinville Limestone, Austinville, VA), 15ml of gypsum (Performance Minerals Corporation, Birmingham, AL), and 39g of 15-9-12 (N-P-K) Osmocote Plus blend (Osmocote, The Scotts Company, Marysville, OH) per pot. Plants were arranged in a split plot design with two treatment ponds per each of 4 plots. Three replicate plants per genotype were placed in each pond. Shallow treatment ponds enabled the bottom 8-10cm of the pots to be submerged in the pond solution. Open water surfaces were covered with black plastic to reduce evaporation and algal growth.

After a week of exposure to fresh water to facilitate establishment, each pond was assigned a salinity treatment of either 0 or 100 mM of sodium chloride (NaCl). All ponds were drained and refilled every day until the treatment ponds reached the target concentration (due to dilution in the soil water). Once treatment concentrations were reached after seven days, ponds were drained and refilled every three days to account for evaporation and maintain a stable concentration within the ponds. To homogenize the distribution of salt within the pots and to prevent salt crystallization on the surface of the soil, pots were top-watered with a ∼200 ml of pond water every time the water was replaced. After treatment initiation, plants were grown for eighteen days and then harvested.

### Measurements and sample collection

At harvest, plants were measured for height and stem diameter and then separated into young leaves (leaves above, and including the most recent fully expanded leaf and bud if present), mature leaves (leaves below the most recent fully expanded leaf, senesced leaves, and cotyledons), stem, and roots. Roots were thoroughly rinsed with fresh water during the root washing. Biomass samples of these four tissue types were dried at 60°C and weighed for dry biomass.

In order to have sufficient tissue for analysis of element content, genotype replicate samples within a pond were pooled for each tissue type, resulting in four biological replicates per tissue type, genotype and treatment. Biomass samples were coarsely ground using a Wiley ^®^ Mini Mill (Thomas Scientific, Swedesboro, NJ) and finely ground with a Qiagen TissueLyser (Qiagen, Germantown, MD) for leaf tissue, and an 8000M Mixer/Mill^®^ High-Energy Ball Mill (SPEX, Metuchen, NJ) for tougher stem and root tissue. Samples were sent to Midwest Laboratories (Midwest Laboratories, Omaha NB) for Inductively Coupled Argon Plasma Optical Emission (ICP) Analysis. Analysis provided the elemental content of boron (B), calcium (Ca), copper (Cu), iron (Fe), potassium (K), magnesium (Mg), manganese (Mn), phosphorus (P), sodium (Na), and sulfur (S), zinc (Zn) for young leaf, mature leaf, stem, and root tissues.

For each biological replicate, the whole plant elemental content was calculated by multiplying the elemental content of each tissue sample by the mass of that tissue, summing all tissue type amounts, and then dividing by the total biomass. The fraction of the element budget allocated to each tissue (e.g. what percentage of total plant sodium was present in the roots) was then calculated from dividing the tissue type element amount by the whole plant element amounts. To compare genotypes that could differ in their organ level element content due to differences in whole plant elemental content we devised a mass allocation based normalization (after (Romero and Maranon, 1996). For each plant we calculated the ratio of the organ mass fraction to the fraction of the element budget in that organ. In this mass relative allocated amount (MRAA) metric, values above one indicate preferential allocation (more of the element is in that tissue then would be based on neutral distribution) and values below one indicate exclusion (less of the element is in that tissue then would be expected based on neutral distribution. To allow for comparison of ratios above and below one we log_2_ transformed the ratios so that a halving and doubling would have the same magnitude (−1, +1). Thus, this MRAA value represents preferential allocation or exclusion of elements from particular tissue types independent of elemental status at the whole plant level. We’ve provided an interactive data sheet that demonstrates the relationship between mass allocation, organ level elemental content, whole plant elemental content and MRAA **(Supplemental datasheet 1)**.

### Data Analysis

All analyses were carried out using R v3.5 (R Foundation for Statistical Computing, Vienna, Austria). For biomass measurements, to combat issues relating to mortality (19 out of 288 plants died) and missing samples (8%, due to labeling error) we averaged tissue types per genotype per treatment pond (taking the mean of natural log transformed weights to minimize skew). This allowed us to construct a complete dataset based on the “average” plant per genotype per treatment pond. Total biomass and biomass fractions of tissue types was then calculated based on the pond average of the tissue types.

Genotype means for all traits in control and stressed environments were then calculated using a mixed-model approach (r package *lme4 (Bates et al., 2015)*) with genotype and treatment as fixed factor and pond within block as random factor. Then from this model estimated marginal means without the random factor were calculated (R package *emmeans*, Lenth, 2018). Significance of main effects and their interaction was determined using Wald’s Chi-square test on the mixed model (R package *car*, Fox and Weisberg, 2011). Using the genotype means per treatment we additionally calculated the change in trait value between control and treatment as the difference in means.

Given the likely correlations between elements we determined common axes of variation using principal component analysis. Principal components were calculated for elemental content at the whole plant level, within each tissue type, and between tissue types at control conditions, salt treatment, and the difference between treatments. Additionally, we calculate the same principal components for MRAA. Differences between tissue types in element content and MRAA were calculated using Hotelling’s-T test on the first two principal components.

Salt tolerance of each genotype was calculated in two ways. First, tolerance was calculated as the proportional change in biomass between both control and salt treatment (taken as the difference between the natural log of total plant biomass in control and salt treatment). Using this metric, genotypes having a smaller negative proportional change have a greater salt tolerance. Then, to follow up on the previous observation that genotypes with high biomass in control conditions have a greater proportional decrease (Temme *et al.*, 2019), a second measure of tolerance was defined as whether a genotype performing better or worse than expected based on size in the control treatment. To quantify this, we fitted a linear regression through the proportional change in biomass vs the natural log of total biomass at control. Using this metric, genotypes with a more positive residual from this linear fit perform better than expected and have a greater salt tolerance.

To determine the effect of trait variation on variation in tolerance we ran a series of linear regressions between tolerance (both metrics) and the first two principal components of whole plant and within tissue type element content and element MRAA at control, salt stressed, and the difference between them. For completeness, all individual component traits of the PC analyses were regressed against tolerance as well. Given the issues of multiple comparisons we limited our interpretation of tolerance associations to the element principal components and only show individual elements as suggestive evidence for an effect on tolerance.

All data visualizations were made using ggplot (Wickham, 2009) and ggbiplot (https://github.com/vqv/ggbiplot)

## Results

### Genotypic differences in salinity tolerance

Genotypes differed in their absolute biomass and response to salinity (**fig 1a, Table 1**), However, proportionally (when comparing log transformed values) the difference between genotypes in the effect of salinity was much more nuanced (**fig 1b**). Across all genotypes we found no evidence for a differential response to salinity. Proportionally all genotypes had a comparable effect of salinity stress. This conclusion is however somewhat simplistic. Although there were no differences among genotypes that rose to the level of significance, there was a significant relationship between vigor (biomass at control conditions) and the proportional effect of salinity (**fig 1c**). Genotypes with a higher vigor had a greater proportional decline in biomass due to salinity stress. Thus, we used two different metrics to characterize tolerance. The first metric was the proportional reduction in biomass: hereafter “proportional-reduction-tolerance.” The second metric took this vigor/salinity effect relationship into account and was the deviation (residual) from the expected effect: hereafter “expectation-deviation-tolerance.”

**Figure 1.**
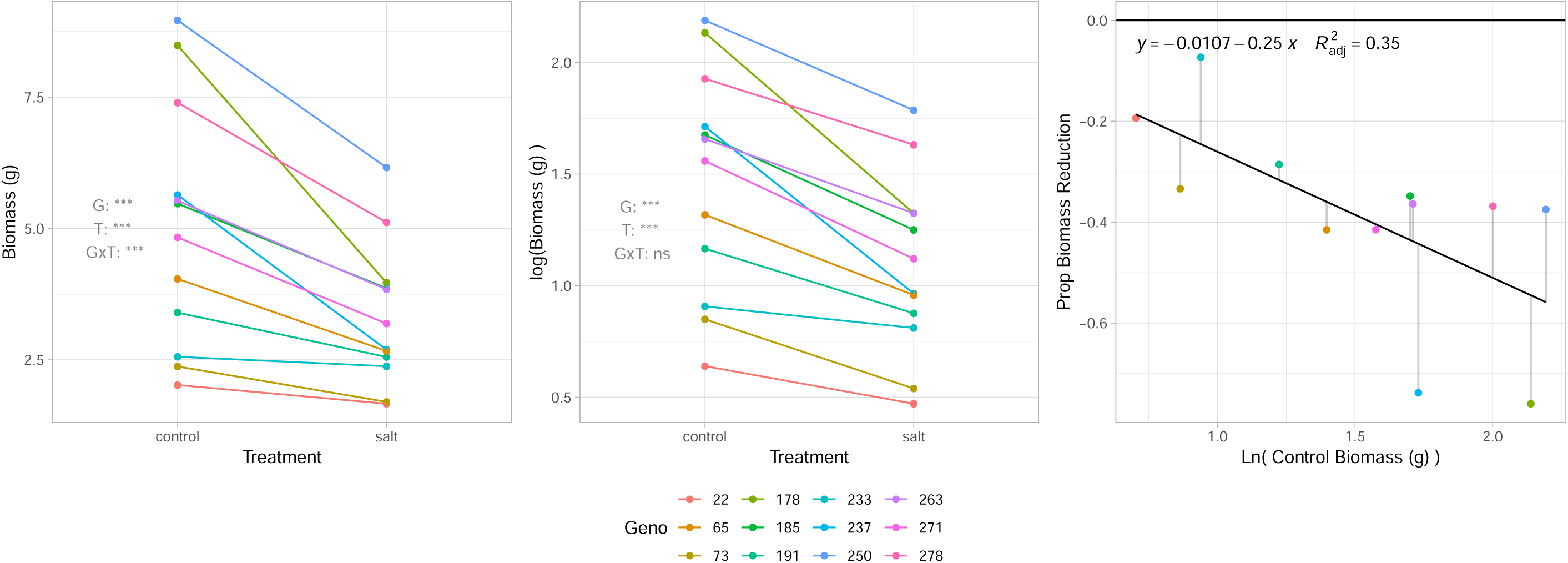
Effect of salinity on biomass and tolerance to salinity. (a) Genotype mean biomass at harvest in control and saline (100 mM NaCl) conditions. (b) Natural log transformed biomass values at control and salt treatment. (c) Relationship between biomass under control conditions and the proportional decline in biomass under salt treatment. Genotypes with more biomass under control conditions have a greater proportional decline in biomass under salt treatment (p<0.05). Different colors denote different genotypes.

**Table 1.** Overview of plant level traits. Median and range (in parentheses) of trait values of 12 genotypes. Stars note p-value significance of Wald’s Chi-square test on treatment (T), genotype (G), and their interaction (TxG). *<0.05, **<0.01, ***<0.001.

| Trait | Control (0mM NaCl) | Salt (100mM NaCl) | T | G | TxG |
| --- | --- | --- | --- | --- | --- |
| *Whole plant mass (g)* | 5.15 (2.02 - 8.96) | 2.94 (1.67 - 6.16) | *** | *** | *** |
| *Young leaf mass (g)* | 1.26 (0.58 - 2.59) | 0.77 (0.47 - 1.85) | *** | *** | ns |
| *Mature leaf mass (g)* | 0.9 (0.37 - 1.82) | 0.58 (0.22 - 1.12) | *** | *** | * |
| *Stem mas (g)* | 1.82 (0.28 - 3) | 0.95 (0.29 - 1.92) | *** | *** | *** |
| *Root mass (g)* | 1 (0.29 - 2.27) | 0.76 (0.39 - 1.44) | ** | *** | ** |
| *YLMF* | 0.26 (0.22 - 0.38) | 0.27 (0.22 - 0.35) | ns | *** | ns |
| *MLMF* | 0.21 (0.17 - 0.27) | 0.19 (0.13 - 0.24) | ns | ns | ns |
| *LMF* | 0.46 (0.4 - 0.64) | 0.45 (0.38 - 0.55) | ns | *** | ns |
| *SMF* | 0.34 (0.11 - 0.4) | 0.28 (0.1 - 0.37) | *** | *** | ns |
| *RMF* | 0.2 (0.12 - 0.27) | 0.23 (0.19 - 0.36) | *** | *** | *** |
| *Height (cm)* | 52.4 (11.36 - 61.32) | 36.57 (11.92 - 43.13) | *** | *** | *** |
| *Stem diameter (mm)* | 8.1 (6.63 - 10.08) | 6.22 (5.3 - 8.88) | *** | *** | ns |
| *SLA (cm<sup>2</sup> g<sup>-1</sup>)* | 368.46 (327.13 - 410.37) | 326.95 (252.61 - 355.63) | *** | *** | ** |
| *B (ppm)* | 7.49 (5.93 - 8.3) | 7.76 (6.84 - 10.34) | *** | *** | * |
| *Ca (%)* | 0.79 (0.65 - 1.09) | 0.7 (0.56 - 0.98) | *** | *** | * |
| *Cu (ppm)* | 2.84 (2.58 - 3.7) | 3.4 (2.82 - 5.1) | *** | *** | ** |
| *Fe (ppm)* | 27.48 (18.92 - 45.39) | 30.6 (23.61 - 54.35) | ** | *** | * |
| *K (%)* | 5.93 (5.54 - 6.82) | 4.58 (3.86 - 5.49) | *** | *** | *** |
| *Mg (%)* | 0.4 (0.36 - 0.58) | 0.33 (0.26 - 0.5) | *** | *** | ns |
| *Mn (ppm)* | 25.25 (20.05 - 37.96) | 34.83 (31.62 - 55.88) | ** | *** | ** |
| *Na (%)* | 0.12 (0.08 - 0.35) | 1.75 (1.21 - 2.59) | *** | *** | *** |
| *P (%)* | 0.4 (0.33 - 0.58) | 0.36 (0.31 - 0.46) | ** | *** | * |
| *S (%)* | 0.53 (0.39 - 0.67) | 0.44 (0.34 - 0.58) | *** | *** | *** |
| *Zn (ppm)* | 15.46 (7.42 - 23.73) | 15.11 (6.15 - 26.83) | ns | *** | ns |

The extremes in the rankings on both metrics lined up. Genotypes with the highest or lowest proportional-reduction-tolerance tended to also be the genotypes with the highest or lowest expectation-deviation-tolerance. However, genotypes in the middle of the ranking differed substantially between tolerance metrics leading to no significant (r_s_ 0.50, *p*=0.099) relationship between tolerance metric ranks (**Fig S1**).

Height and stem diameter, traits strongly related to biomass showed comparable results with biomass (**Table 1, Figure S2**). Allocation of mass to different plant organs (young leaf, mature leaf, stem, and root) showed a mixed response with no effect of treatment on leaf mass fraction (LMF) and no interaction between genotype and treatment on LMF and stem mass fraction. Root mass fraction did differ significantly between genotypes and treatment including a significant interaction. While not a trait of focus we did measure specific leaf area (SLA) which had a significant effect of treatment, genotype and an interaction between them. (**Table 1, Figure S2**)

### Whole plant and organ level elements

Depending on element, genotypes differed in their whole plant elemental content and in the effect of salinity thereon. (**Table 1, Figure S3**) Given the highly correlated nature of elemental content, we sought to capture the variation in elemental content in principal component analysis.

At the whole plant level principal component analysis revealed that under control and salt treatment over 70% of the variation in elemental content could be captured by the first two principal components (**Fig 2a,b**). Results showed strong, negative, relationships between whole plant sodium and potassium content and between boron and zinc content. Depending on treatment there were strong positive relationships between elements though these shifted between treatments. Additionally, over 50% of the variation in the response of whole plant elemental content, assessed as the difference between control and salt treatment, was captured in the first two principal components. The positive and negative relationships among shifts in elemental content were similar to those of elemental content itself (**Fig 2c**).

**Figure 2.**
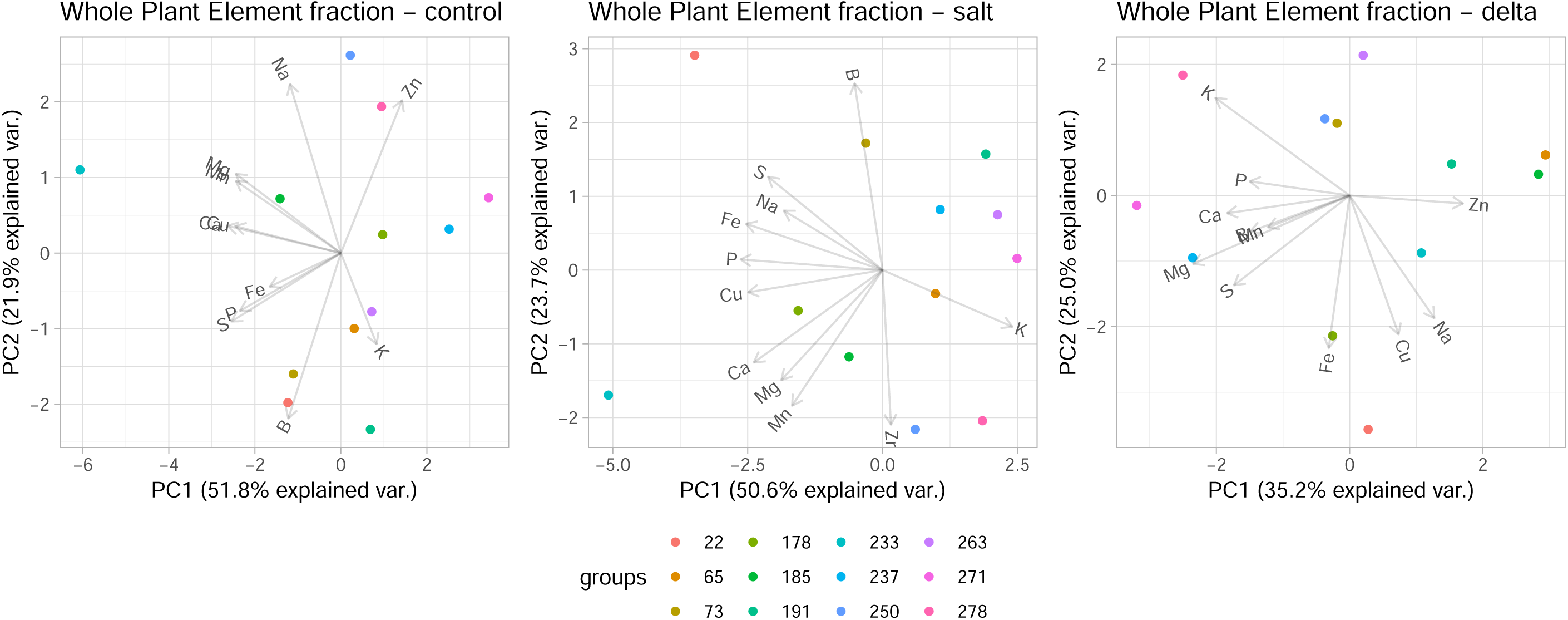
Multivariate view of whole plant elemental content. First and second principal components of whole plant elemental content under (a) control treatment, (b) salt treatment, and (c) the difference between treatments. Under both treatments >70% of the variability in elemental content for 11 macro and micro nutrients was captured in the first two principal components. >60% of the variability in the change of elemental content was captured in the first two principal components. Different colors denote different genotypes.

Within organs, over 60% in the variation in element content and MRAA under saline conditions could be captured in the first two principal components (**Figure S4, S5**). Contrasting individual plant organs, again, depending on the element, genotypes largely differed in their organ level elemental content (**Table 2**) and in the mass relative allocated amount (MRAA)(**Table 3**) to their organs (**Figure S6**). Notably, under salt stress, sodium content increased the most in stems and roots and the least in leaves. Potassium content showed the opposite pattern. (**Table 2**). In terms of mass relative allocated amount (MRAA), genotypes were capable of keeping the majority of sodium constrained to the roots and out of the leaves under control conditions **(Table 3, Figure S3)**. However, under salt stress the MRAA in leaves became less negative indicating sodium inevitably reached these sensitive tissues. Due to this the level of preferential allocation to the roots also decreased. Surprisingly, MRAA in stems greatly increased, going from nearly neutral to positive showing increased allocation of sodium to stems under saline conditions **(Table 3, Figure S3)**. Potassium MRAA showed an opposite pattern.

**Table 2.** Organ level elemental content. Median and range (in parentheses) of trait values of 12 genotypes. Stars note p-value significance of Wald’s Chi-square test on treatment (T), genotype (G), organ (O) and their interactions. .<0.1, *<0.05, **<0.01, ***<0.001. Letters note significant Tukey post hoc groups based on contrasting all eight organ/treatment groups.

| Element | Treatment | Young Leaves | Mature Leaves | Stem | Root | T | G | O | GxT | GxO | TxO | GxTxO |
| --- | --- | --- | --- | --- | --- | --- | --- | --- | --- | --- | --- | --- |
| B (ppm) | control | 9.31 (7.47 - 14.88) <sup>a</sup> | 15.53 (12.5 - 18.35) <sup>c</sup> | 2.47 (2.17 - 3.12) <sup>e</sup> | 3.04 (2.45 - 3.33) <sup>e</sup> | *** | *** | *** | ns | *** | *** | ns |
|  | salt | 10.9 (8.53 - 15.58) <sup>b</sup> | 17.68 (14.83 - 20.03) <sup>d</sup> | 2.8 (2.15 - 4.67) <sup>e</sup> | 2.86 (2.32 - 3.1) <sup>e</sup> |  |  |  |  |  |  |  |
| Ca (%) | control | 0.88 (0.74 - 1.16) <sup>a</sup> | 1.12 (0.76 - 1.79) <sup>c</sup> | 0.59 (0.52 - 0.72) <sup>d</sup> | 0.57 (0.4 - 0.61) <sup>e</sup> | *** | *** | *** | *** | *** | *** | *** |
|  | salt | 0.82 (0.61 - 0.99) <sup>b</sup> | 1.12 (0.77 - 1.8) <sup>c</sup> | 0.47 (0.4 - 0.52) <sup>e</sup> | 0.51 (0.42 - 0.66) <sup>de</sup> |  |  |  |  |  |  |  |
| Cu (ppm) | control | 2.41 (2.05 - 2.6) <sup>ab</sup> | 2.03 (1.78 - 2.52) <sup>bc</sup> | 1.23 (0.65 - 2.77) <sup>d</sup> | 7.28 (6.2 - 9.38) <sup>e</sup> | *** | *** | *** | ** | *** | ns | * |
|  | salt | 2.81 (2.02 - 3.5) <sup>a</sup> | 2.72 (1.86 - 3.4) <sup>a</sup> | 1.26 (0.95 - 3.17) <sup>cd</sup> | 8.15 (5.92 - 9.89) <sup>f</sup> |  |  |  |  |  |  |  |
| Fe (ppm) | control | 11.07 (8.9 - 70.53) <sup>a</sup> | 11.74 (9.57 - 32.02) <sup>a</sup> | 5.31 (3.1 - 12.67) <sup>a</sup> | 87.76 (61.95 - 169.98) <sup>b</sup> | ns | *** | *** | ** | *** | ** | *** |
|  | salt | 10.27 (6.8 - 13.35) <sup>a</sup> | 12.42 (7.92 - 35) <sup>a</sup> | 7.31 (3.83 - 15.02) <sup>a</sup> | 101.86 (80.33 - 149.54) <sup>b</sup> |  |  |  |  |  |  |  |
| K (%) | control | 5.32 (5 - 6.34) <sup>a</sup> | 7.37 (6.17 - 8.77) <sup>b</sup> | 6.94 (5.45 - 8.5) <sup>c</sup> | 3.37 (3.02 - 3.99) <sup>e</sup> | *** | *** | *** | *** | *** | *** | *** |
|  | salt | 5.47 (4.83 - 6.27) <sup>a</sup> | 7.32 (4.98 - 9.24) <sup>bc</sup> | 4.62 (3.24 - 5.02) <sup>d</sup> | 2.05 (1.49 - 2.51) <sup>f</sup> |  |  |  |  |  |  |  |
| Mg (%) | control | 0.38 (0.31 - 0.58) <sup>ab</sup> | 0.4 (0.28 - 0.9) <sup>cd</sup> | 0.53 (0.46 - 0.66) <sup>e</sup> | 0.18 (0.14 - 0.2) <sup>f</sup> | *** | *** | *** | * | *** | *** | ** |
|  | salt | 0.38 (0.28 - 0.58) <sup>ac</sup> | 0.47 (0.33 - 0.94) <sup>d</sup> | 0.33 (0.23 - 0.43) <sup>b</sup> | 0.17 (0.13 - 0.19) <sup>f</sup> |  |  |  |  |  |  |  |
| Mn (ppm) | control | 29.94 (23.58 - 41.2) <sup>ab</sup> | 37.4 (25.7 - 59.25) <sup>cd</sup> | 19.35 (15.35 - 23.83) <sup>f</sup> | 20.7 (13.15 - 30.9) <sup>f</sup> | ** | *** | *** | *** | *** | *** | *** |
|  | salt | 39.62 (32.5 - 61.07) <sup>ac</sup> | 49.4 (37.1 - 103.07) <sup>e</sup> | 26.79 (22.45 - 31.73) <sup>bdf</sup> | 26.86 (22.9 - 38.47) <sup>bdf</sup> |  |  |  |  |  |  |  |
| Na (%) | control | 0.01 (0 - 0.19) <sup>a</sup> | 0.01 (0 - 0.02) <sup>a</sup> | 0.12 (0.05 - 0.79) <sup>ab</sup> | 0.43 (0.28 - 0.93) <sup>b</sup> | *** | *** | *** | *** | *** | *** | *** |
|  | salt | 0.42 (0.11 - 1.25) <sup>b</sup> | 1.01 (0.37 - 1.84) <sup>c</sup> | 2.96 (1.71 - 5.75) <sup>d</sup> | 2.58 (1.81 - 2.78) <sup>e</sup> |  |  |  |  |  |  |  |
| P (%) | control | 0.61 (0.5 - 0.78) <sup>a</sup> | 0.34 (0.28 - 0.53) <sup>c</sup> | 0.25 (0.2 - 0.45) <sup>d</sup> | 0.42 (0.31 - 0.6) <sup>e</sup> | ** | *** | *** | *** | *** | *** | ns |
|  | salt | 0.52 (0.38 - 0.57) <sup>b</sup> | 0.29 (0.24 - 0.35) <sup>d</sup> | 0.24 (0.2 - 0.38) <sup>d</sup> | 0.4 (0.31 - 0.51) <sup>ce</sup> |  |  |  |  |  |  |  |
| S (%) | control | 0.64 (0.6 - 0.77) <sup>a</sup> | 0.73 (0.62 - 0.94) <sup>c</sup> | 0.26 (0.14 - 0.39) <sup>d</sup> | 0.51 (0.41 - 0.69) <sup>b</sup> | *** | *** | *** | *** | *** | *** | ns |
|  | salt | 0.5 (0.42 - 0.57) <sup>b</sup> | 0.65 (0.46 - 0.85) <sup>a</sup> | 0.22 (0.17 - 0.29) <sup>d</sup> | 0.46 (0.36 - 0.62) <sup>b</sup> |  |  |  |  |  |  |  |
| Zn (ppm) | control | 5.68 (4.65 - 8.47) <sup>a</sup> | 18.46 (12.25 - 33.37) <sup>bc</sup> | 8.77 (2.67 - 36.42) <sup>b</sup> | 26.16 (5.7 - 64.67) <sup>cd</sup> | ns | *** | *** | ns | *** | * | *** |
|  | salt | 5.5 (4.54 - 7.88) <sup>a</sup> | 14.1 (9.17 - 33.73) <sup>b</sup> | 8.54 (2.04 - 43.06) <sup>b</sup> | 35.23 (5.63 - 57.93) <sup>d</sup> |  |  |  |  |  |  |  |

**Table 3.** Element mass relative allocated amount (MRAA) per organ. Median and range (in parentheses) of trait values of 12 genotypes. Stars note p-value significance of Wald’s Chi-square test on treatment (T), genotype (G), organ (O) and their interactions. .<0.1, *<0.05, **<0.01, ***<0.001. Letters note significant Tukey post hoc groups based on contrasting all eight organ/treatment groups.

| Element | Treatment | Young Leaves | Mature Leaves | Stem | Root | T | G | O | GxT | GxO | TxO | GxTxO |
| --- | --- | --- | --- | --- | --- | --- | --- | --- | --- | --- | --- | --- |
| B | control | 0.36 (0.06 - 0.74) <sup>a</sup> | 1.18 (0.74 - 1.26) <sup>b</sup> | -1.5 (-1.79 - -1.27) <sup>c</sup> | -1.28 (-1.49 - -0.98) <sup>d</sup> | ns | * | *** | ** | *** | *** | ns |
|  | salt | 0.48 (-0.02 - 0.67) <sup>a</sup> | 1.23 (0.72 - 1.37) <sup>b</sup> | -1.53 (-1.88 - -1.18) <sup>c</sup> | -1.48 (-1.75 - -1.27) <sup>c</sup> |  |  |  |  |  |  |  |
| Ca | control | 0.2 (0.01 - 0.43) <sup>a</sup> | 0.52 (0.18 - 0.71) <sup>b</sup> | -0.36 (-1.01 - -0.09) <sup>d</sup> | -0.51 (-1.05 - -0.23) <sup>e</sup> | ns | *** | *** | ns | *** | *** | *** |
|  | salt | 0.2 (-0.08 - 0.42) <sup>a</sup> | 0.72 (0.41 - 0.88) <sup>c</sup> | -0.53 (-1.08 - -0.37) <sup>e</sup> | -0.45 (-0.83 - -0.18) <sup>d</sup> |  |  |  |  |  |  |  |
| Cu | control | -0.28 (-0.76 - -0.04) <sup>a</sup> | -0.58 (-0.95 - -0.13) <sup>b</sup> | -1.24 (-2 - -0.44) <sup>c</sup> | 1.27 (0.98 - 1.51) <sup>d</sup> | *** | *** | *** | ** | *** | *** | * |
|  | salt | -0.41 (-1 - 0.06) <sup>ab</sup> | -0.37 (-0.98 - 0.02) <sup>ab</sup> | -1.46 (-1.82 - -0.7) <sup>c</sup> | 1.13 (0.82 - 1.32) <sup>e</sup> |  |  |  |  |  |  |  |
| Fe | control | -1.11 (-1.98 - 0.14) <sup>a</sup> | -1.03 (-1.87 - -0.5) <sup>a</sup> | -2.41 (-3.36 - -1.54) <sup>c</sup> | 1.7 (0.82 - 2.11) <sup>d</sup> | ** | *** | *** | *** | *** | *** | *** |
|  | salt | -1.81 (-2.13 - -1.17) <sup>b</sup> | -1.11 (-2.22 - -0.79) <sup>a</sup> | -2.21 (-3.04 - -1.17) <sup>c</sup> | 1.66 (1.18 - 1.85) <sup>d</sup> |  |  |  |  |  |  |  |
| K | control | -0.15 (-0.31 - 0.09) <sup>a</sup> | 0.31 (0.11 - 0.5) <sup>b</sup> | 0.24 (-0.08 - 0.47) <sup>b</sup> | -0.81 (-0.96 - -0.64) <sup>d</sup> | * | *** | *** | . | *** | *** | *** |
|  | salt | 0.23 (0.06 - 0.49) <sup>b</sup> | 0.62 (0.29 - 0.76) <sup>c</sup> | -0.11 (-0.45 - 0.19) <sup>a</sup> | -1.17 (-1.64 - -0.83) <sup>e</sup> |  |  |  |  |  |  |  |
| Mg | control | -0.08 (-0.35 - 0.09) <sup>a</sup> | 0.07 (-0.39 - 0.64) <sup>c</sup> | 0.38 (0.18 - 0.75) <sup>d</sup> | -1.17 (-1.67 - -0.9) <sup>e</sup> | *** | *** | *** | ns | *** | *** | *** |
|  | salt | 0.22 (0.02 - 0.37) <sup>b</sup> | 0.47 (0.02 - 0.91) <sup>d</sup> | -0.02 (-0.66 - 0.48) <sup>ac</sup> | -1.04 (-1.55 - -0.68) <sup>f</sup> |  |  |  |  |  |  |  |
| Mn | control | 0.26 (-0.2 - 0.56) <sup>a</sup> | 0.53 (0.27 - 0.66) <sup>b</sup> | -0.44 (-0.94 - -0.13) <sup>c</sup> | -0.38 (-0.99 - -0.05) <sup>c</sup> | ns | *** | *** | ns | *** | ns | *** |
|  | salt | 0.21 (-0.12 - 0.45) <sup>a</sup> | 0.56 (0.07 - 0.87) <sup>b</sup> | -0.48 (-0.81 - -0.21) <sup>c</sup> | -0.37 (-0.84 - -0.05) <sup>c</sup> |  |  |  |  |  |  |  |
| Na | control | -4.09 (-5.56 - -2.61) <sup>a</sup> | -4.17 (-5.87 - -2.7) <sup>a</sup> | -0.18 (-1.26 - 0.39) <sup>d</sup> | 1.58 (1.13 - 2.04) <sup>f</sup> | *** | *** | *** | * | *** | *** | ns |
|  | salt | -2.38 (-3.8 - -0.74) <sup>b</sup> | -1.11 (-2.47 - 0.03) <sup>c</sup> | 0.72 (-0.51 - 1.71) <sup>e</sup> | 0.5 (-0.48 - 1.16) <sup>e</sup> |  |  |  |  |  |  |  |
| P | control | 0.57 (0.42 - 0.77) <sup>a</sup> | -0.16 (-0.44 - 0.18) <sup>c</sup> | -0.63 (-0.98 - -0.41) <sup>e</sup> | 0.05 (-0.3 - 0.33) <sup>f</sup> | ns | * | *** | ns | *** | *** | ** |
|  | salt | 0.43 (0.14 - 0.72) <sup>b</sup> | -0.33 (-0.58 - -0.12) <sup>d</sup> | -0.63 (-0.74 - -0.19) <sup>e</sup> | 0.15 (-0.09 - 0.31) <sup>f</sup> |  |  |  |  |  |  |  |
| S | control | 0.33 (-0.1 - 0.66) <sup>a</sup> | 0.55 (0.26 - 0.78) <sup>c</sup> | -1.04 (-1.53 - -0.52) <sup>d</sup> | -0.06 (-0.19 - 0.37) <sup>e</sup> | ns | *** | *** | ns | *** | *** | * |
|  | salt | 0.16 (-0.17 - 0.54) <sup>b</sup> | 0.55 (0.32 - 0.83) <sup>c</sup> | -0.99 (-1.28 - -0.57) <sup>d</sup> | 0.01 (-0.15 - 0.31) <sup>e</sup> |  |  |  |  |  |  |  |
| Zn | control | -1.45 (-1.81 - -0.28) <sup>a</sup> | 0.08 (-0.44 - 1.67) <sup>bc</sup> | -0.7 (-1.54 - 0.61) <sup>d</sup> | 0.27 (-0.61 - 1.65) <sup>bc</sup> | ns | ns | *** | ns | *** | * | *** |
|  | salt | -1.36 (-2.12 - -0.1) <sup>a</sup> | -0.17 (-1.23 - 1.66) <sup>b</sup> | -0.75 (-1.38 - 0.66) <sup>d</sup> | 0.97 (-0.91 - 1.63) <sup>c</sup> |  |  |  |  |  |  |  |

While genotypes differed in the elemental content and MRAA in a given organ, the difference among organs was far greater than the difference among genotypes. Over 70% of the variation in elemental content between organs could be captured in the first two principal components with organs having a significantly different suite of elemental content. Under benign conditions, genotypes had a greater sodium, copper and iron content in roots and more manganese, boron and calcium in leaves (both young and mature) (**Figure 3a)**. Under saline conditions, this changed such that stems and then roots had a greater increase in sodium content and that leaves had a lower decrease in potassium content (**Figure 3b,c**). Separation of tissue types on element MRAA was comparable to element content (**Figure 3d-f**).

**Figure 3.**
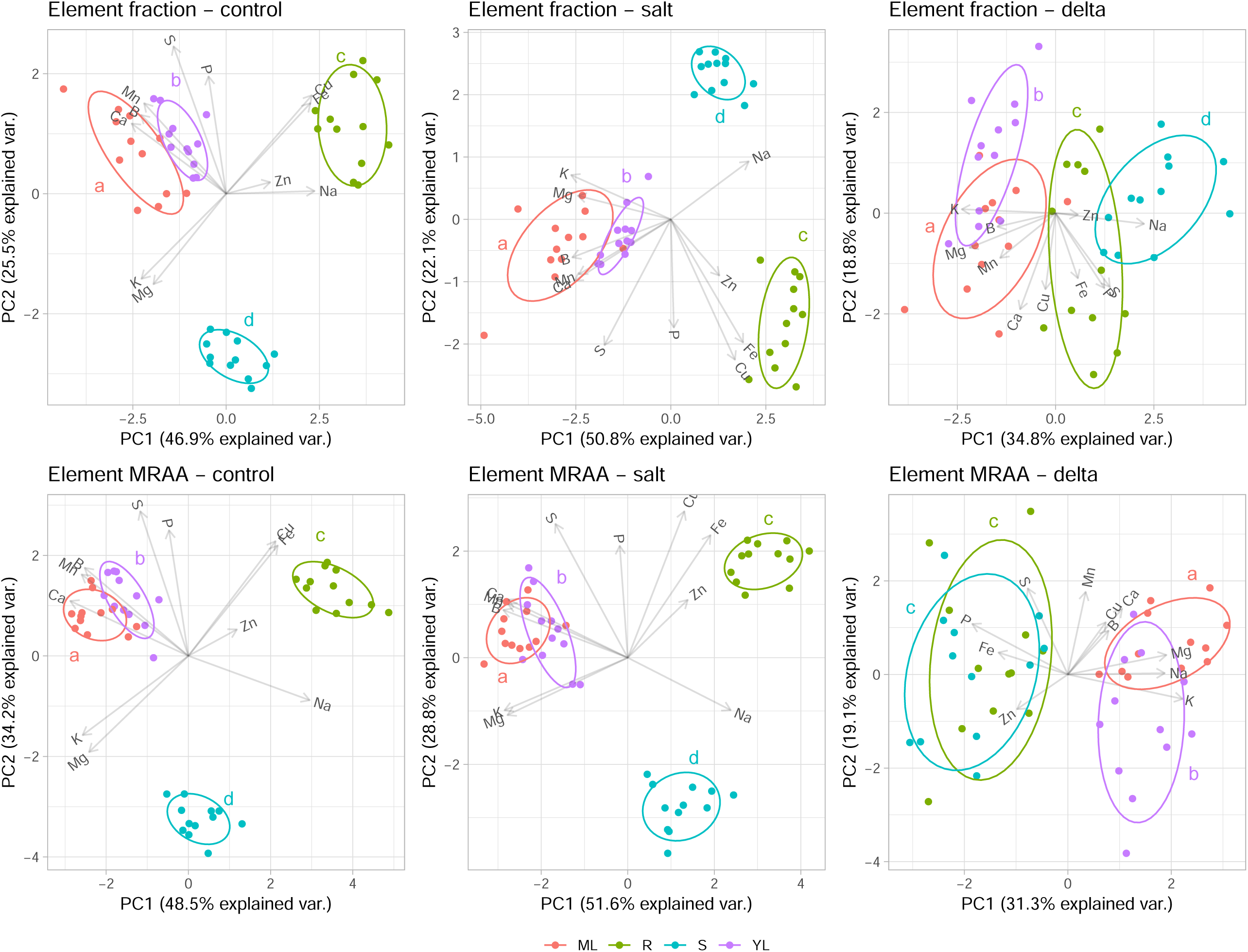
Multivariate view of organ level elemental content. First and second principal component of organ level elemental content under (a) control treatment, (b) salt treatment, and (c) the difference between treatments. Organ types, leaf (young and mature), stem, and root, were significantly differentiated with the first two PC axes capturing >70% of the variability between tissues. Letters denote Hottelings-T significant differences on the first two PC axes.

### Relationships between mass allocation, elements and salinity tolerance

We performed a series of linear regressions to determine the association between elemental traits and tolerance metrics. Given the large number of comparisons this brings up the issue of false discoveries. Thus, while we present these results, care should be taken in interpreting causative mechanisms from weakly significant results. To partially combat this here we focus on the relationship between the principal components of elemental content and allocation with tolerance instead of each individual element. It must be noted however that even these relationships do not hold up to stringent multiple comparisons correction and should only be interpreted as indicative of potential associations.

At the whole plant level, elemental content was not strongly associated with tolerance (**Table S1**). Only the first principal component of elemental content in control conditions (**Fig 2a**) was associated with proportional-reduction-tolerance. The elements most strongly tied to this axis were calcium, sulfur, and copper (**Table 4**). This result likely recapitulates the vigor/effect of salinity relationship since this same PC axis is suggestively correlated with vigor under control conditions (p<0.1).

**Table 4.** Element loadings on PCA axes linked with tolerance. Fraction of variation in element content or MRAA that loads onto the PCA axes that suggestively (non-significant after multiple comparison correction) associate with variation in tolerance (*Table S4*). Rank of fraction explained in parentheses. WP: whole plant, M leaf: mature leaf,

| Element | Proportional-reduction-tolerance |  |  | Expectation-deviation-tolerance |  |
| --- | --- | --- | --- | --- | --- |
|  | WP content<br>PC1 (control) | Stem content<br>PC1 (salt) | M leaf MRAA<br>PC1 (delta) | Root content<br>PC2 (salt) | Root content<br>PC2 (delta) |
| B | 0.19 (9) | 0.30 (7) | **0.61 (1)** | **0.29 (3)** | 0.02 (9) |
| Ca | **0.88 (1)** | >0.01 (10) | **0.53 (3)** | 0.01 (10) | 0.06 (8) |
| Cu | **0.76 (3)** | 0.60 (4) | 0.43 (5) | 0.05 (6) | 0.06 (7) |
| Fe | 0.34 (7) | 0.01 (9) | >0.01 (11) | 0.06 (5) | 0.36 (4) |
| K | 0.09 (11) | 0.50 (6) | 0.02 (9) | **0.75 (1)** | **0.44 (2)** |
| Mg | 0.75 (4) | >0.01 (11) | 0.17 (8) | 0.23 (4) | 0.12 (6) |
| Mn | 0.75 (5) | **0.78 (1)** | 0.53 (4) | 0.05 (7) | 0.02 (10) |
| Na | 0.18 (10) | 0.53 (5) | 0.40 (6) | **0.42 (2)** | **0.60 (1)** |
| P | 0.69 (6) | **0.73 (2)** | **0.55 (2)** | 0.02 (9) | >0.01 (11) |
| S | **0.81 (2)** | **0.63 (3)** | 0.01 (10) | 0.05 (8) | **0.38 (3)** |
| Zn | 0.25 (8) | 0.14 (8) | 0.19 (7) | >0.01 (11) | 0.17 (5) |

As a contributor to variation in MRAA we related changes in biomass allocation to tolerance as well. Biomass allocation to plant organs was differentially associated with both metrics of tolerance **(Table S2**. Proportional-reduction-tolerance was negatively correlated with leaf mass fraction at control conditions. Although this simply recapitulates the correlation found in fig 1c due to the fact that genotypes with more biomass had a lower leaf mass fraction. Expectation-deviation-tolerance was correlated with the change in stem mass fraction (genotypes with a lower reduction in stem mass fraction had a lower than expected decline).

When we related within organ principal components (**Figure S4, S5**) to both metrics of tolerance (**Table S3**) a contrasting picture emerged. Proportional-reduction-tolerance was associated with the first principal component of stem elemental content under saline conditions (**Figure 4a)** (with highest contribution to manganese, phosphorus and sulfur content (**Table 4**)) as well as the second principal component of the change in MRAA of elements to mature leaves (**Figure 4b)** (with highest contribution to boron, phosphorous, and calcium MRAA change (**Table 4**)). Expectation-deviation-tolerance was associated with the second principal component of root elemental content under saline conditions (**Figure 4c**)(highest contribution to potassium, sodium and boron (**Table 4**)) as well as second principal component of the change in root elemental content (**Figure 4d**) (highest contributions to sodium, potassium and sulfur (**Table 4**)).

**Figure 4.**
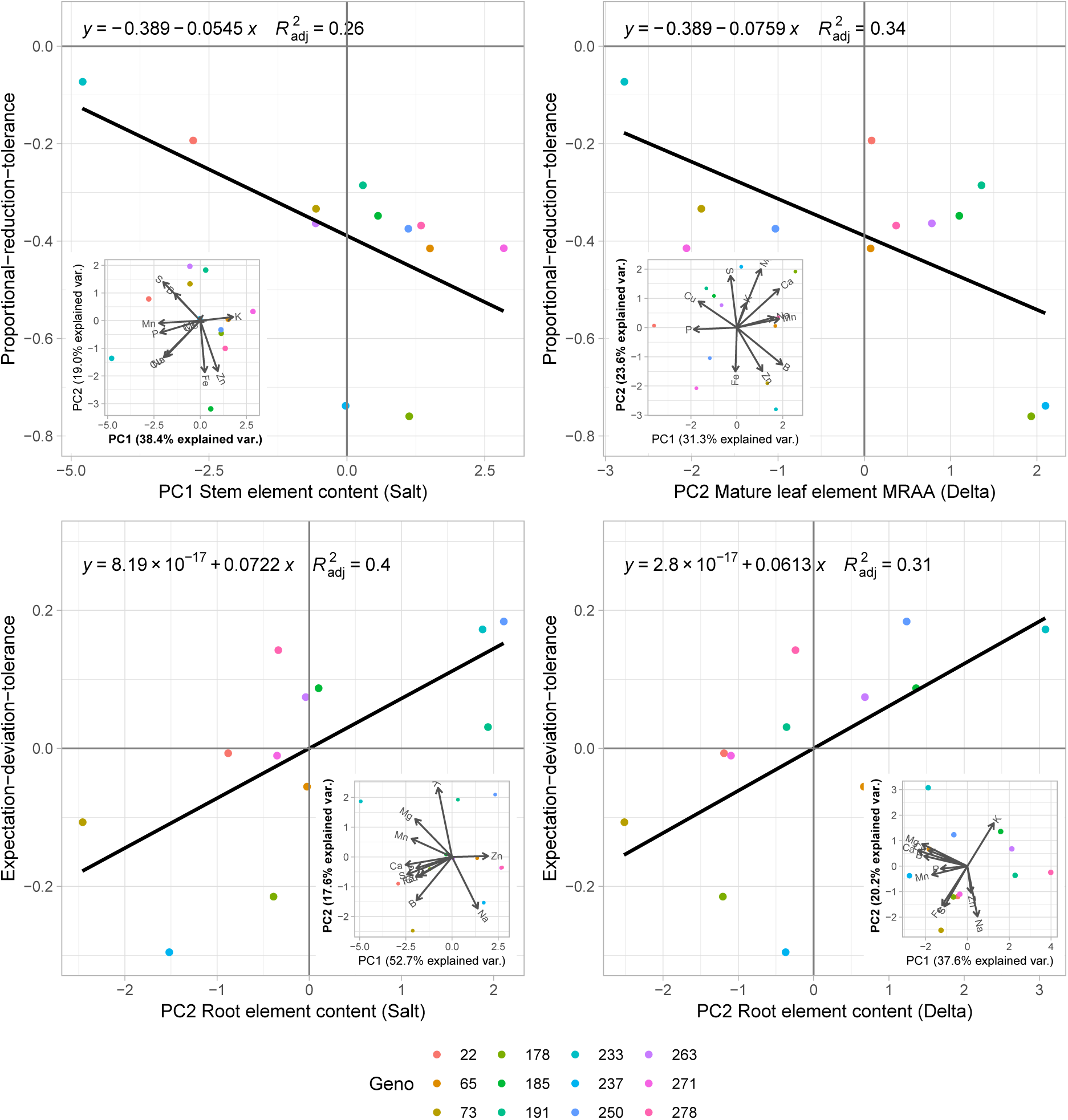
Relationship between “significant” organ level elemental content principal components and tolerance. Proportional-reduction tolerance and (a) stem element content under salt treatment PC1 and (b) the change in mature leaf element MRAA PC2. Expectation-deviation-tolerance and (c) root element content under salt treatment PC2 and (d) the change in root element content PC2. Panel insets show the PCA graph corresponding to the panel. Note that while all slopes are p<0.05, these are non-significant after multiple comparisons correction (Table 4).

Surprisingly, the elements that contributed the most to the principal components associated with either tolerance metric weren’t necessarily the most significantly associated with tolerance when looked at all elements individually. We found putatively significant relationships between element content, MRAA and both metrics of tolerance but the sheer number of multiple comparisons (>100) lead to these being weakly suggestive at best. **(Figure S7, Table S4**)

## Discussion

Salinity is a major abiotic stress decreasing crop yield by limiting water uptake and disrupting cell metabolism and ion homeostasis. Mitigating some of the adverse effects requires some combination of energy intensive sodium exclusion or sequestration in vacuoles and/or non-photosynthesizing tissues (Tyerman *et al.*, 2019; Munns *et al.*, 2019). Indeed, in cultivated sunflower prior work has shown a trade-off where genotypes more vigorous in benign conditions are proportionally more affected by salinity than less vigorous genotypes (Temme *et al.*, 2019). High tolerance could come at the cost of vigor under benign conditions (Munns and Gilliham, 2015). However, here we show that tolerance can be redefined by explicitly taking this trade-off into account by scoring genotypes on their deviation from the expected proportional decline. This expectation-deviation-tolerance allows us to identify different traits associated with genotype performing better or worse than expected and potentially allows for the selection of varieties that break away from this vigor/effect of salinity trade-off.

Across twelve cultivated sunflower genotypes, we replicated the vigor/proportional decline in biomass trade-off (Temme *et al.*, 2019). It should be noted that while this relationship is significant and explains a moderate portion of the variance in proportional decline (**Fig 1c**, p<0.05, R^2^_adj_ 0.35) the magnitude of variation in proportional decline between genotypes is such that it doesn’t rise to the level of significance when simply contrasting genotypes (**Fig 1b**). When contrasting genotypes on two metrics of tolerance, one based solely on the proportional decline, proportional-decline-tolerance, and one on deviation from the expected proportional decline based on a genotypes vigor, expectation-deviation-tolerance, we found genotypes ranking to be uncorrelated (**Fig S7**). Given this discrepancy in ranking we further explored the link between variation in genotypes elemental traits and the connection with both tolerance metrics.

Element content and allocation was assessed for eleven elements (Na, P, K, Ca, Mg, S, Fe, B, Mn, Cu, Zn) in both control and saline conditions as well as the change between them. We found surprisingly strong correlations between elemental content at the whole plant level (**Fig 2a-b**) and clear differentiation in elemental content between organs (**Table 2, Fig 3a-b**). At the whole plant level, under both benign and stressed conditions, there were strong negative correlations between sodium and potassium. This likely reflects variation in uptake selectivity and the antagonistic nature of sodium and potassium uptake (Blumwald, 2000; Mäser *et al.*, 2002). Additionally, there was a similar negative relationship between boron and zinc which have a negative relationship in maize (Hosseini *et al.*, 2007) and wheat (Singh *et al.*, 1990). Out of eleven elements, only whole plant zinc content was unaffected by salinity (**Table 1, Fig S1**). These results show the strong effects of high soil salinity on plant elemental content beyond those typically measured (generally Na and K (Mäser *et al.*, 2002; Wu *et al.*, 2018).

We further investigated the impact of salinity stress on the content and distribution of elements to leaves (young and mature), stem, and roots. Prior research has shown elemental distribution to be element specific and impacted by stress (Conn and Gilliham, 2010). Within organs elemental content was strongly correlated as well with differences between organs being far greater than differences between genotypes (**Fig 3a-b**). Our results support the notion that in multivariate space, variation between tissue types (even between young and mature leaves) is far greater than the variation between genotypes (Agren and Weih, 2012). Besides elemental content, we investigated the effect of salinity on the preferential allocation of elements to tissue types. In this mass relative allocated amount (MRAA), we assessed the deviation from a neutral distribution by calculating the fold change of the proportion of mass allocated to an organ and the proportion of whole plant elemental amount allocated to the organ (after (Romero and Maranon, 1996). Given the predominant role of sodium and potassium, we highlight these elements.

For sodium, this revealed that under benign conditions plants preferentially keep sodium out of the leaves and keep it in the roots (**Table 3, Fig S3**). Under salinity stress, mechanisms maintaining Na into the root appear unable to cope and Na content rises throughout the plant **(Table 2, Fig S3**). Mass relative allocation changes as well with less exclusion of sodium from the leaves and relatively less preferential allocation to the roots. Interestingly, MRAA of sodium to the stem increases (**Table 3, Fig S3**). These results indicate that mechanisms pulling sodium out of the transpiration stream at the root level (liken to HKT1 transporters (Munns and Tester, 2008; Hasegawa, 2013)) are overwhelmed but that mechanisms at the stem or leaf level kick into gear. Possible mechanisms could include high activity of tonoplast transporters sequestering Na into the vacuole in stem tissue (Bassil *et al.*, 2019), transporters pulling out sodium in the petiole (Jeschke and Pate, 1991), or active phloem loading, moving Na out of the leaf (Zhong *et al.*, 1998).

Potassium distribution broadly mirrors sodium with elevated MRAA of potassium to leaves and lower MRAA in roots under benign conditions. Under saline conditions, this pattern is magnified with even greater potassium MRAA to leaves and less to roots (**Table 3, Fig S3**). As at the whole plant level potassium content decreases, this heightened differentiation between plant organs showcases the key role of potassium in leaf processes ((Hawkesford *et al.*, 2012). In the stems, the greater accumulation and allocation of sodium is not matched by an increase in potassium MRAA. Since potassium and sodium have antagonistic effects in the cytosol maintaining a proper balance between these ions is vital (Munns and Tester, 2008). The discrepancy between stem sodium and potassium content and allocation would thus suggest sequestration in the vacuole to be a possible mechanism of sodium accumulation here to maintain cytosolic Na:K ratio’s while elevating Na sequestration in the stem tissue (Blumwald, 2000; Zeng *et al.*, 2018; Bassil *et al.*, 2019).

Despite the effects of salinity stress on whole plant elemental content, the mass relative allocated amount (MRAA) was not dramatically affected for elements other than sodium or potassium. Interestingly, copper, iron, sodium and zinc have a positive MRAA in roots (both under control and stressed conditions) whereas for all other elements it’s neutral or negative. This is opposite findings in wheat (Garnett and Graham, 2005) and carrot (Inal *et al.*, 2009). It could be that this reflects ions bound to the root apoplast from the soil solution (Negrão *et al.*, 2017) but given our thorough washing and the lower MRAA of other elements in roots, that seems not to be a major factor. Given sunflower’s status as a moderate heavy metal accumulator (Adesodun *et al.*, 2010), this points to active exclusion of these elements from the shoot or a functional role in the root for these elements.

Linking variation in elemental traits to variation in tolerance has some inherent difficulties in our experimental design. Given the large number of traits versus a comparatively small number of genotypes, multiple comparisons corrections are a large dampener on the significance of our findings. Care must be taken in interpreting our findings, as chance playing a role in our findings cannot be ruled out. For future work testing hypotheses generated by our results we strongly recommend a larger number of genotypes to reach greater power to detect links between a continuum of tolerance values and a host of elemental traits.

Bearing in mind multiple comparisons issues on significance, we found that the associations between tolerance and elemental traits differed by tolerance metric. Associated with proportional-decline-tolerance were shifts in mature leaf element allocation and stem elemental content under salt stress. Associated with expectation-deviation-tolerance were root elemental content and it’s shifts under salt stress. (**Fig 4**) Top elements loading onto these multivariate axes were manganese and boron for proportional-reduction-tolerance and potassium and sodium for expectation-deviation-tolerance (**Table 4**).

Strikingly, these top loading elements were not necessarily the elements that were associated with tolerance when looked at individually. Though fraught with multiple comparisons correction issues (>100 comparisons), when looking at each individual element in each plant part in turn, results support the role of root potassium content for expectation-deviation-tolerance but also suggest a role for root and plant sulfur as well as leaf and plant manganese content. Sulfur content has been previously found to play a role in sunflower salt tolerance (Temme *et al.*, 2019). Root sulpholipid content has been linked to salinity tolerance (Erdei *et al.*, 1980). Maintenance of manganese content is linked to salt tolerance in brassica (Chakraborty *et al.*, 2016). Manganese is tied to salinity stress in it’s component of manganese superoxide dismutase which functions as a reactive oxygen species (ROS) scavenger and the oxidative component of salt stress (Hernández *et al.*, 1994; Wang *et al.*, 2010).

Given the large number of traits increasing the number of genotypes assessed for these elemental traits will shed more light on significant association with salt tolerance. Moreover, given the short duration of this experiment a next step will be to track element status across developmental stage and it’s impact on salinity tolerance beyond the seedling stage. Developmental status has a large impact on element distribution in wheat (Weih *et al.*, 2016), barley (Birsin *et al.*, 2010). With the strong trade-off between plant vigor (biomass in control conditions) and proportional-reduction-tolerance and the limited overlap between traits connected with both metrics of tolerance, these results suggest the possibility for decoupling vigor related traits and tolerance. By decoupling vigor from tolerance, we could select for high yielding genotypes that are able to maintain their growth under saline conditions. A key open question then is whether these tolerance and vigor traits share a common genetic basis or if tolerance related genes could be pyramided into high vigor varieties (Morton *et al.*, 2018).

Here we found that increased sodium sequestration in stems and roots and increased potassium allocation to leaves appears to be a common response due to salinity. However, given the highly correlated nature of organ elemental content, it is difficult to pinpoint whether variation in these specific traits underpins variation in tolerance. What is clear is that the multivariate nature of tolerance associated traits needs to be considered especially for elemental content. Our results suggest that taking vigor related trade-offs into account can be vital in determining the traits related to tolerance. By selecting on traits that allow a genotype to perform better than expected, we could potentially break free from these trade-offs and maintain high vigor in a saline environment.

## Supporting information

Element allocation (MRAA) example sheet

## Acknowledgements

We would like to thank the University of Georgia greenhouse staff and several undergraduate students for their aid in set up and harvest of the experiment. This work was financially supported by grant NSF1444522 to LAD.

## Figure captions

**Figue S1.** Genotypes ranking on tolerance metrics. Each genotype’s rank on expectation-deviation-tolerance versus proportional-reduction-tolerance. Spearman’s correlation of ranks is non-significant (r_s_=0.50, p=0.099), dashed line notes 1:1 line.

**Figure S2.** Phenotypic traits measured at harvest. Stem Diam (stem diameter), CCI (chlorophyll content index), Plant height, Specific Leaf Area (SLA), Young Leaf Mass Fraction (YLMF), Mature Leaf Mass Fraction (MLMF), Leaf Mass Fraction (LMF), Stem Mass Fraction (SMF), and Root Mass Fraction (RMF). Different colors note different genotypes. Insets show genotype, treatment and their interaction p-value (Wald’s chi-square). .<0.1, *<0.05, **<0.01, ***<0.001.

**Figure S3.** Whole elemental content reaction norms between control and salt (100mM NaCl) treatment. Different colors note different genotypes. Insets show genotype, treatment and their interaction p-value (Wald’s chi-square). .<0.1, *<0.05, **<0.01, ***<0.001.

**Figure S4.** Within organ multivariate view of elemental content. First and second principal components of per organ elemental content under salt treatment and of the difference between control and salt treatment. Different colors denote different genotypes.

**Figure S5.** Within organ multivariate view of element mass relative allocated amount (MRAA). MRAA shows the log2 fold difference between the proportion of element allocated to an organ and the proportion of mass allocated to an organ. First and second principal components of per organ MRAA under salt treatment and of the difference between control and salt treatment. Different colors denote different genotypes.

**Figure S6.** Organ specific reaction norms of elemental content. Set of figures (one per element) displaying organ level (Young Leaf, Mature Leaf, Stem, Root) reaction norms of elemental content (top row), element distribution (middle row), and Mass Relative Allocated Amount (MRAA, bottom row). MRAA shows the log2 fold difference between the proportion of element allocated to an organ and the proportion of mass allocated to an organ. Different colors note different genotypes. Insets show genotype, treatment and their interaction p-value (Wald’s chi-square). .<0.1, *<0.05, **<0.01, ***<0.001.

**Figure S7.** Tolerance and element correlation. Multi page supplement on the correlation between tolerance, defined as the residuals of the log biomass/proportional decrease in biomass relationship and organ level elemental content under salt treatment (page 1), the change in elemental content (page 2), and the proportional decrease in biomass and organ level elemental content under salt treatment (page 3), the change in elemental content (page 4). Individual elements are arranged in rows with organ classes in columns. Young leaf (YL), mature leaf (ML), stem (S), Root (R), and whole plant (Total).

## Notes

#### Summary of Updates

added supplementary files

