## Supplementary tables for "Element content and distribution has limited, tolerance metric dependent, impact on salinity tolerance in cultivated sunflower (*Helianthus annuus*)"

**Table S1.** Association between whole plant elemental content principal components and tolerance. P-value significance and model R2 of the correlation of the first and second principal component of whole plant elemental content under control, salt treatment and the change between them and tolerance where tolerance is defined as either the proportional decrease in biomass or the residuals of the proportional decrease/vigor relationship.

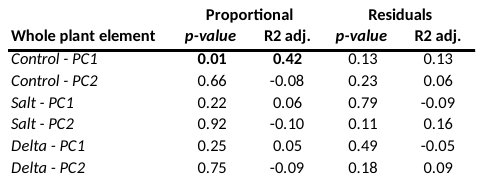

**Table S2.** Association between plant morphology and tolerance. P-value significance and model R2 of the correlation of organ level mass fractions under control and salt treatment and the change between them and tolerance where tolerance is defined as either the proportional decrease in biomass or the residuals of the proportional decrease/vigor relationship. Young leaf (YL), mature leaf (ML), stem (S), Root (R).

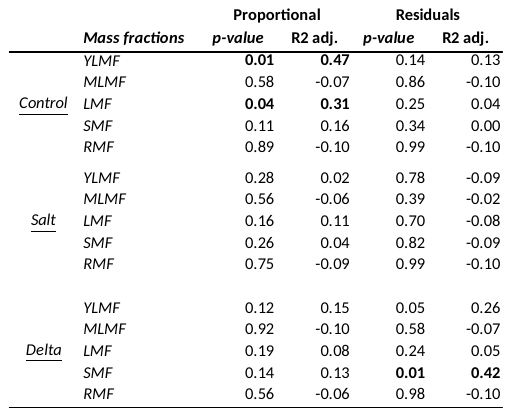

**Table S3.** Association between tissue level elemental content and mass relative allocated amount principal components and tolerance. P-value significance and model R2 of the correlation of the first and second principal component of organ level plant elemental content and MRAA under salt treatment and the change between them and tolerance where tolerance is defined as either the proportional decrease in biomass or the residuals of the proportional decrease/vigor relationship. Young leaf (YL), mature leaf (ML), stem (S), Root (R).

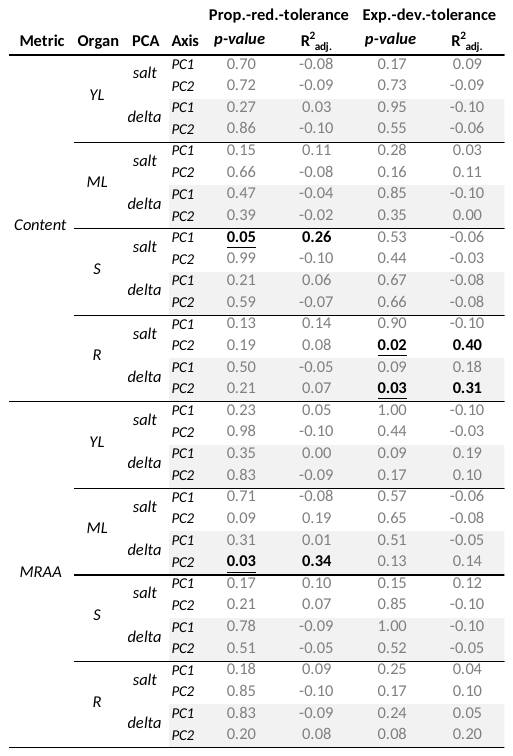

**Table S4.** Association between organ (and whole plant) level elemental content, elemental MRAA and both metrics of tolerance. P-value significance and model R2 of individual element metric and tolerance. P fdr notes Benjamini-Hochberg false discovery rate multiple comparison correction. Green shaded pvalues <0.05.

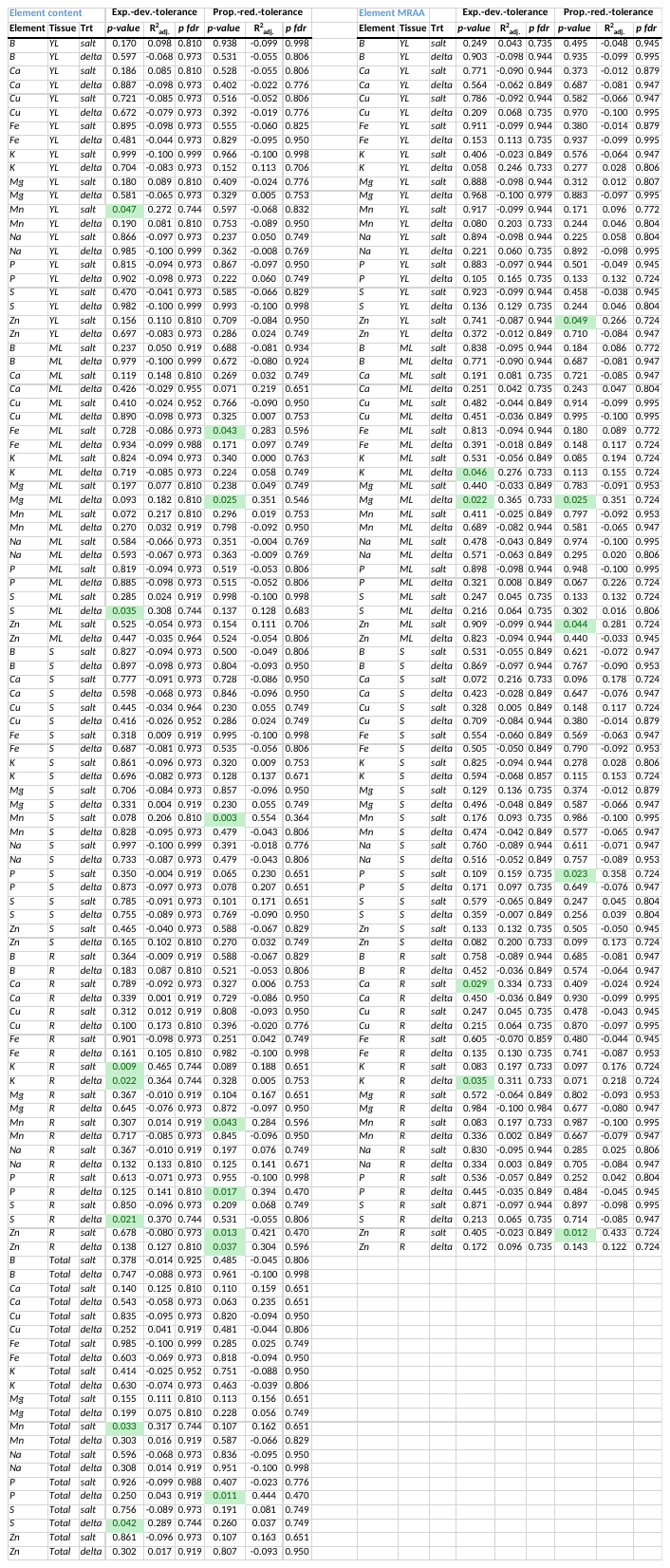
