## Supplementary figures and images for "Element content and distribution has limited, tolerance metric dependent, impact on salinity tolerance in cultivated sunflower (*Helianthus annuus*)"

### Figure S1

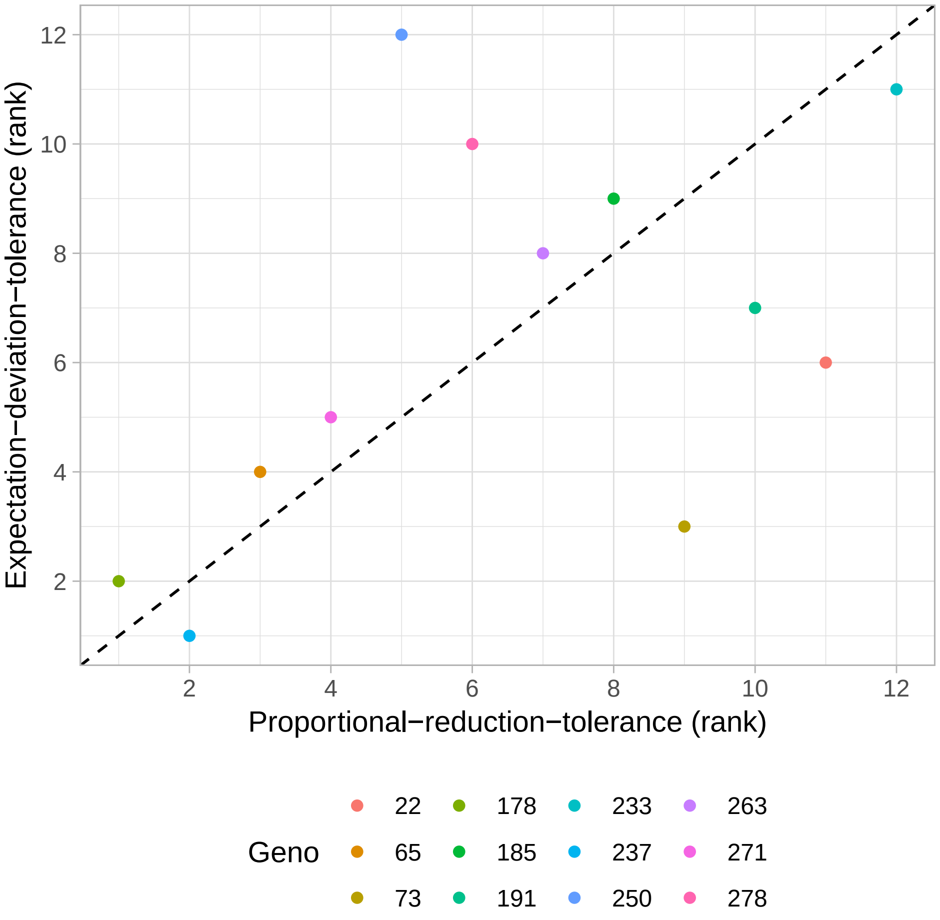

### Figure S2

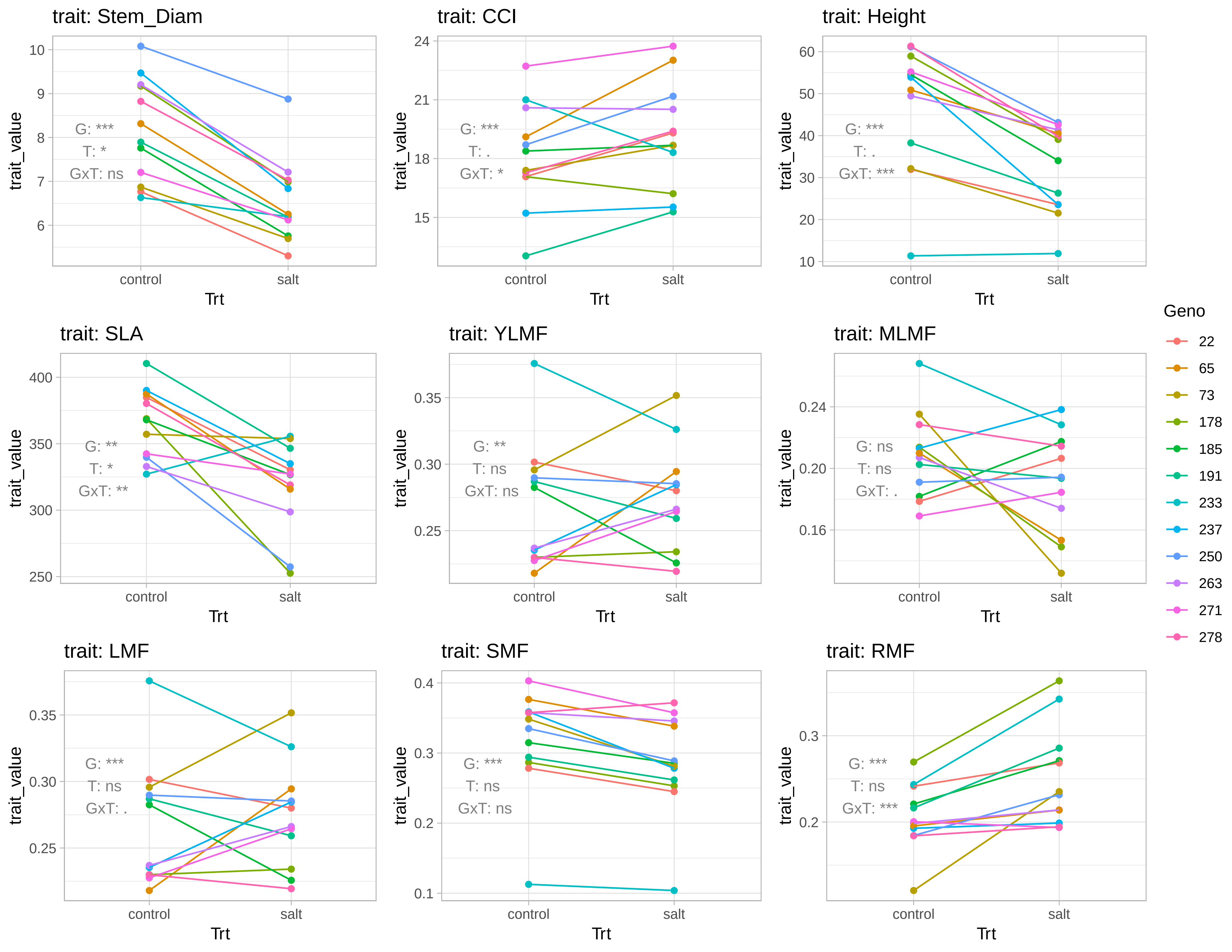

### Figure S3

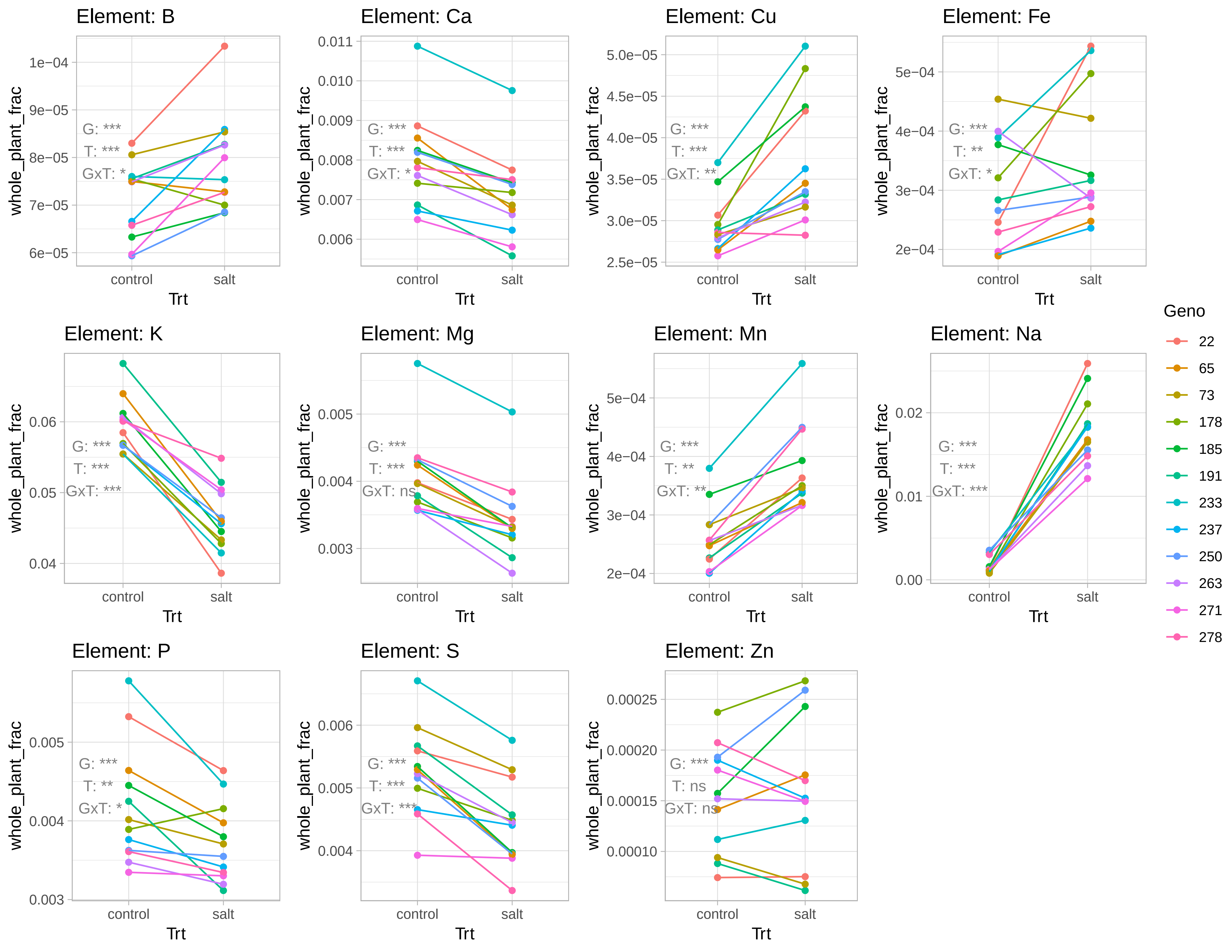

### Figure S4

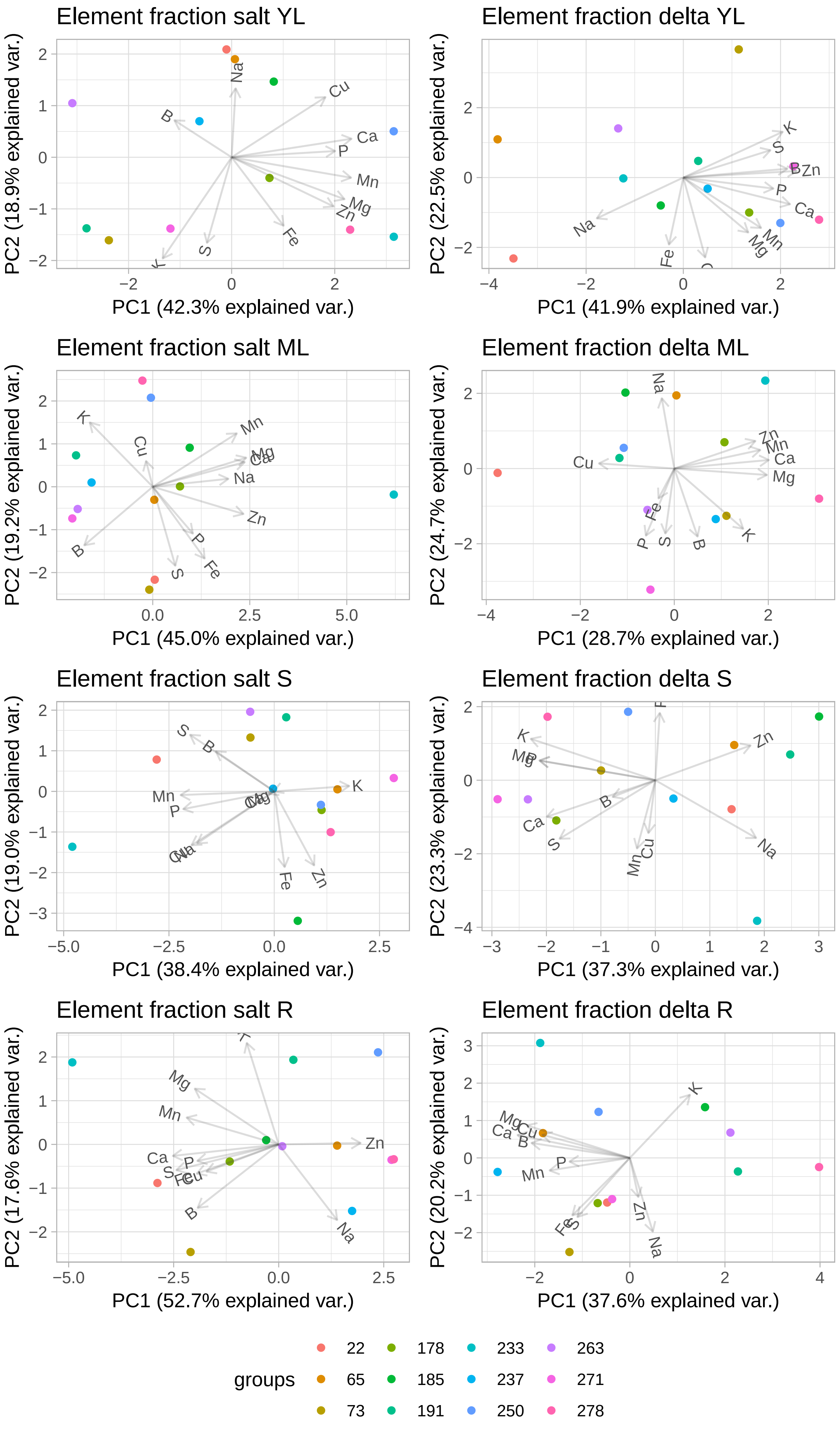

### Figure S5

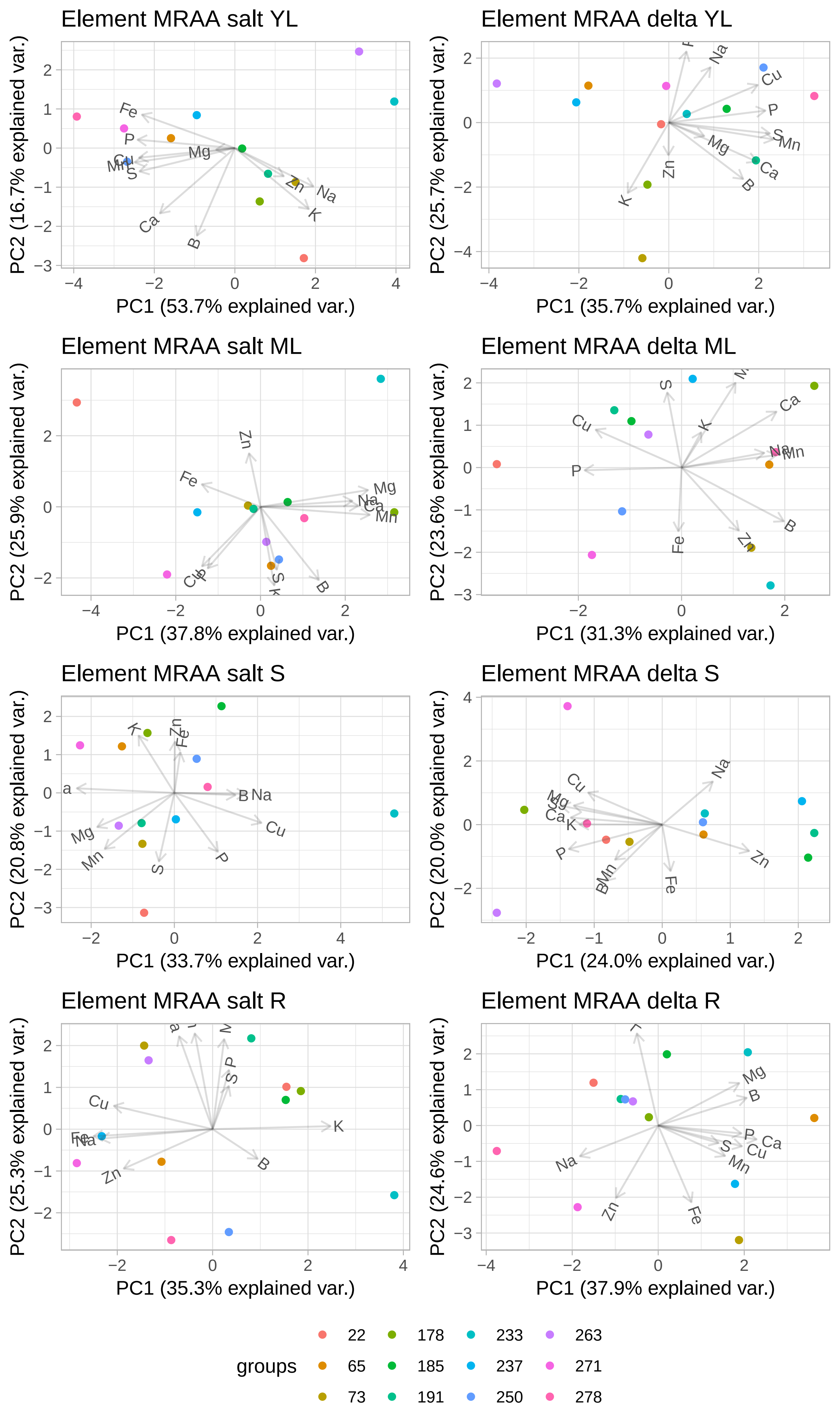

### Figure S7

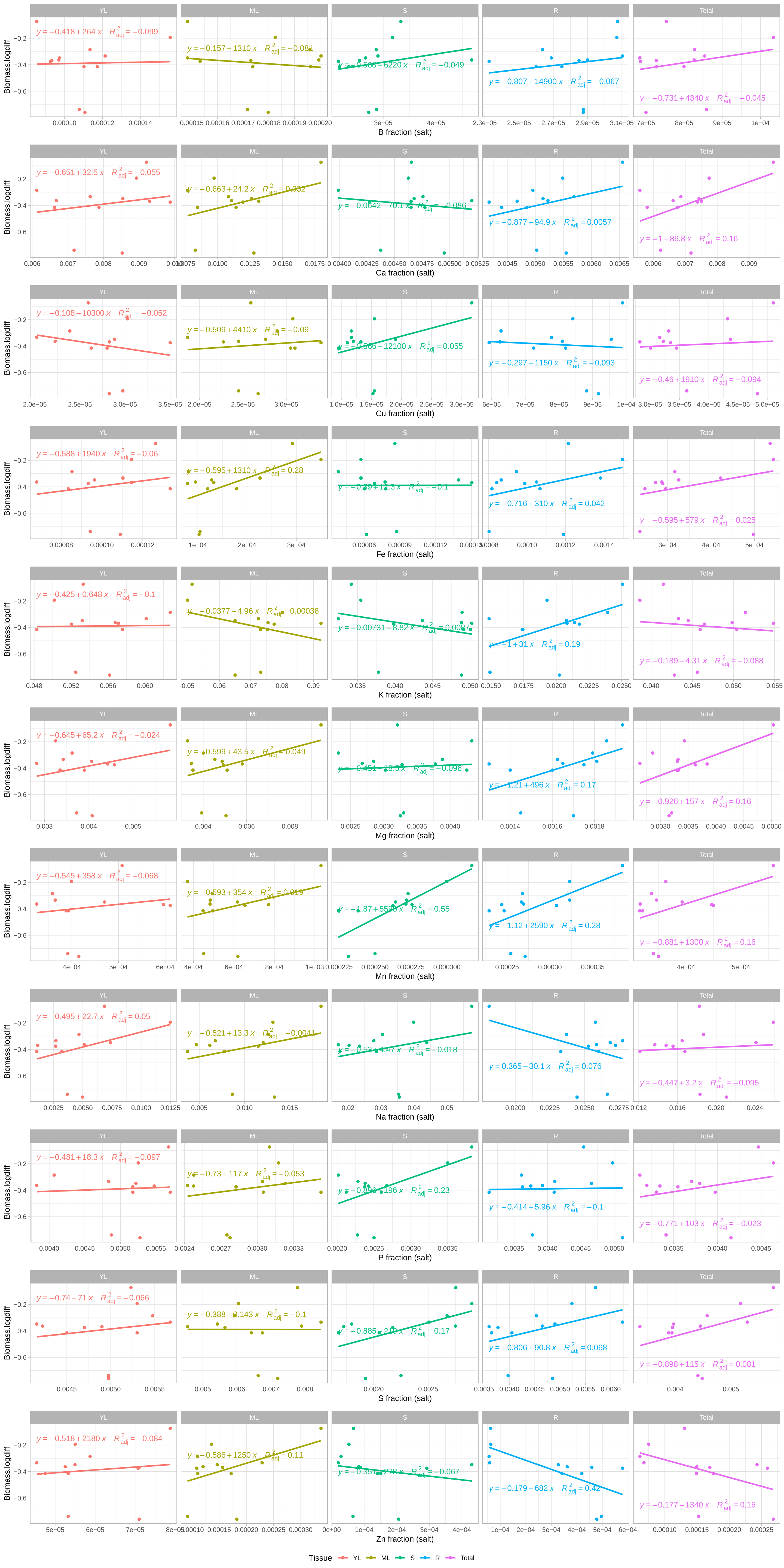

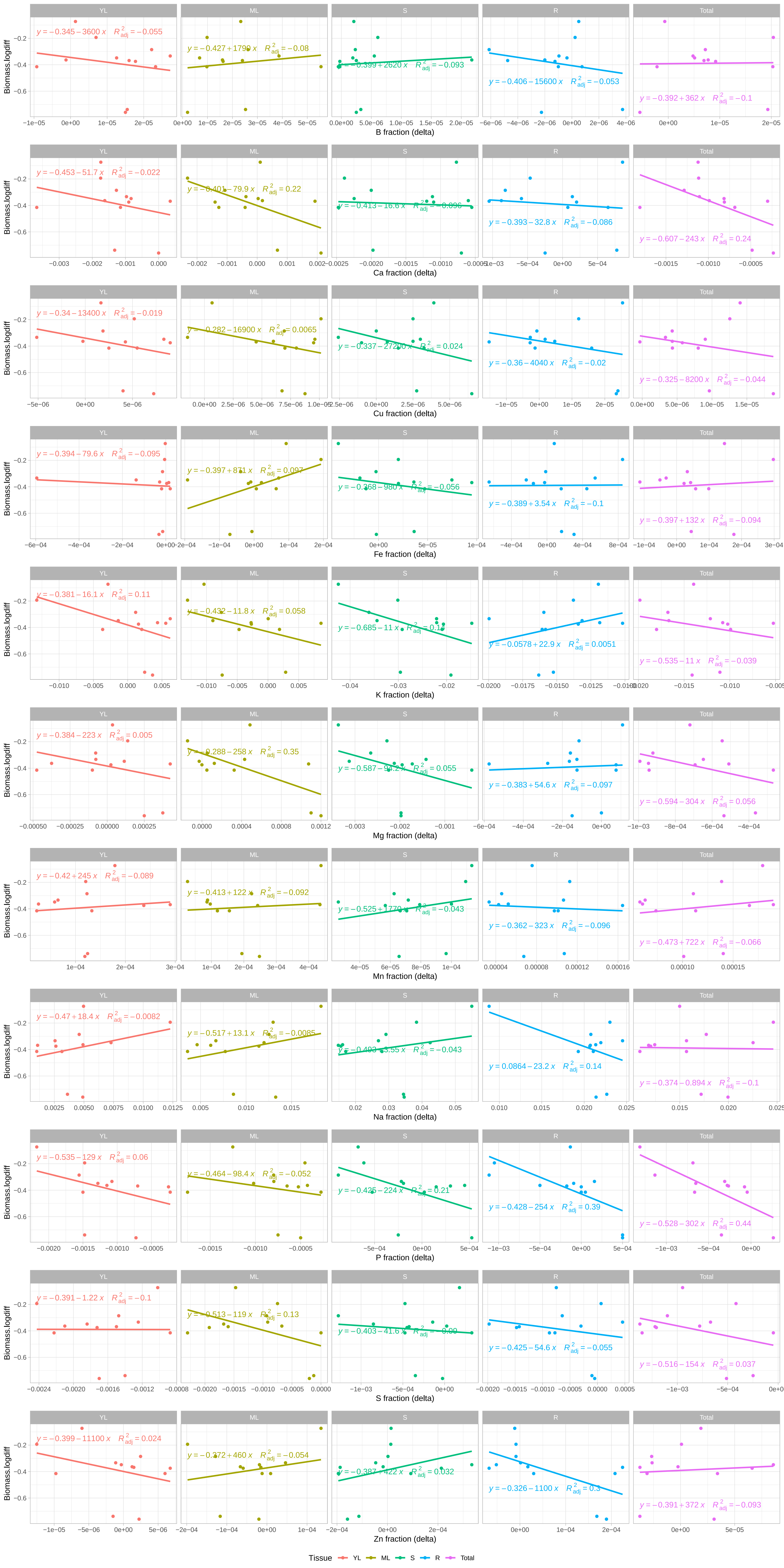

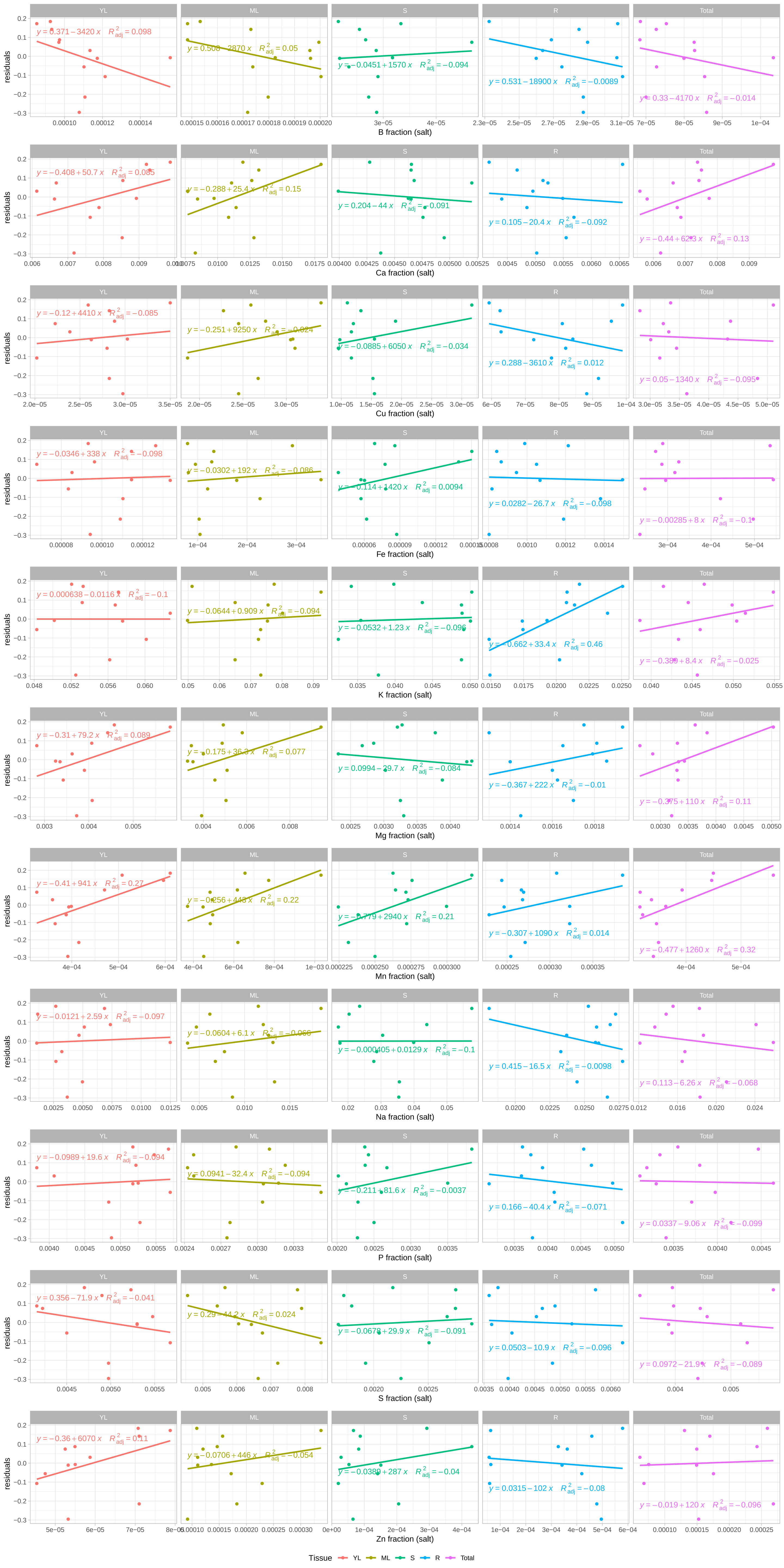

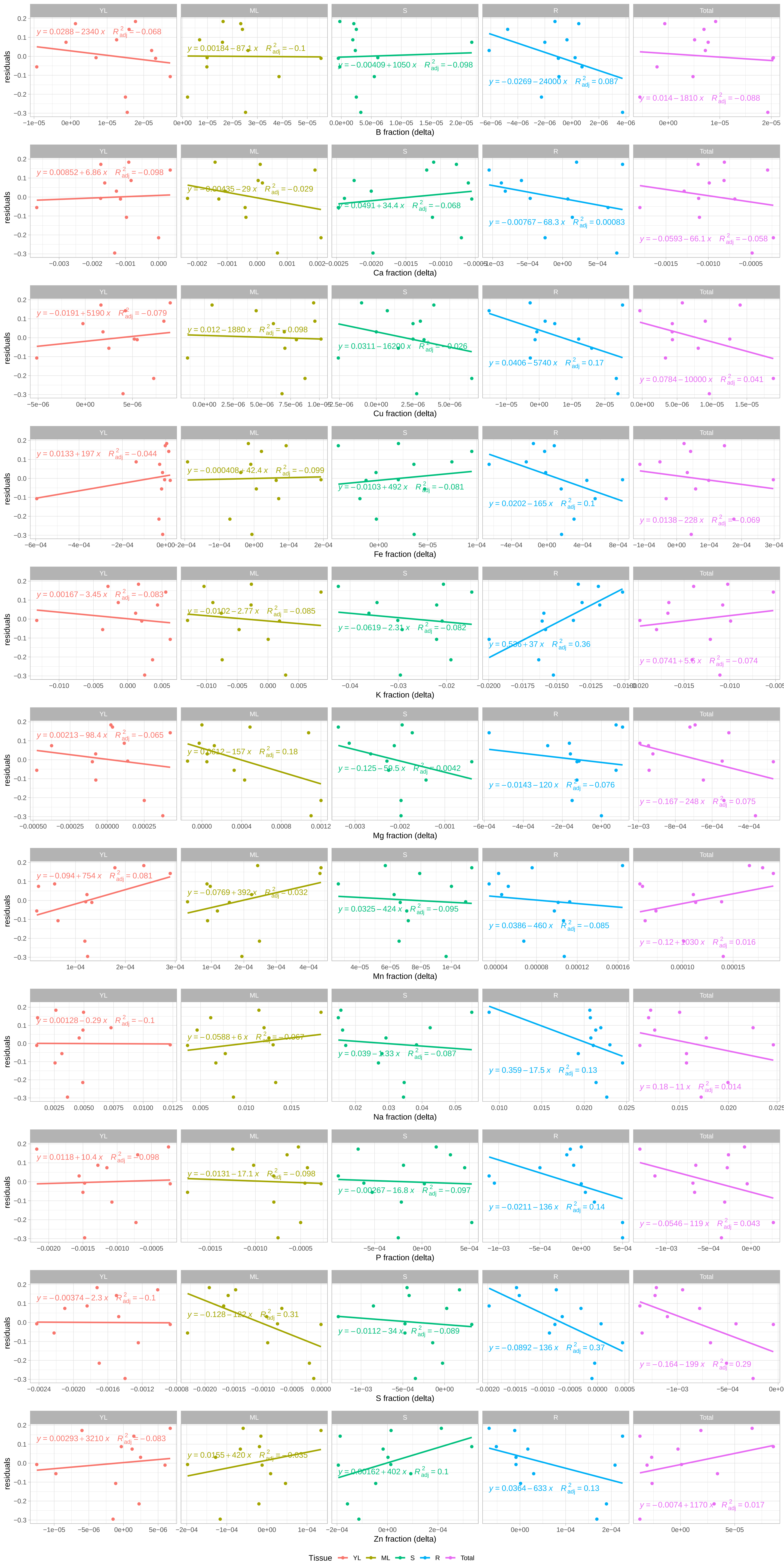
