## Supplementary material for "Element content and distribution has limited, tolerance metric dependent, impact on salinity tolerance in cultivated sunflower (*Helianthus annuus*)": Figure S6

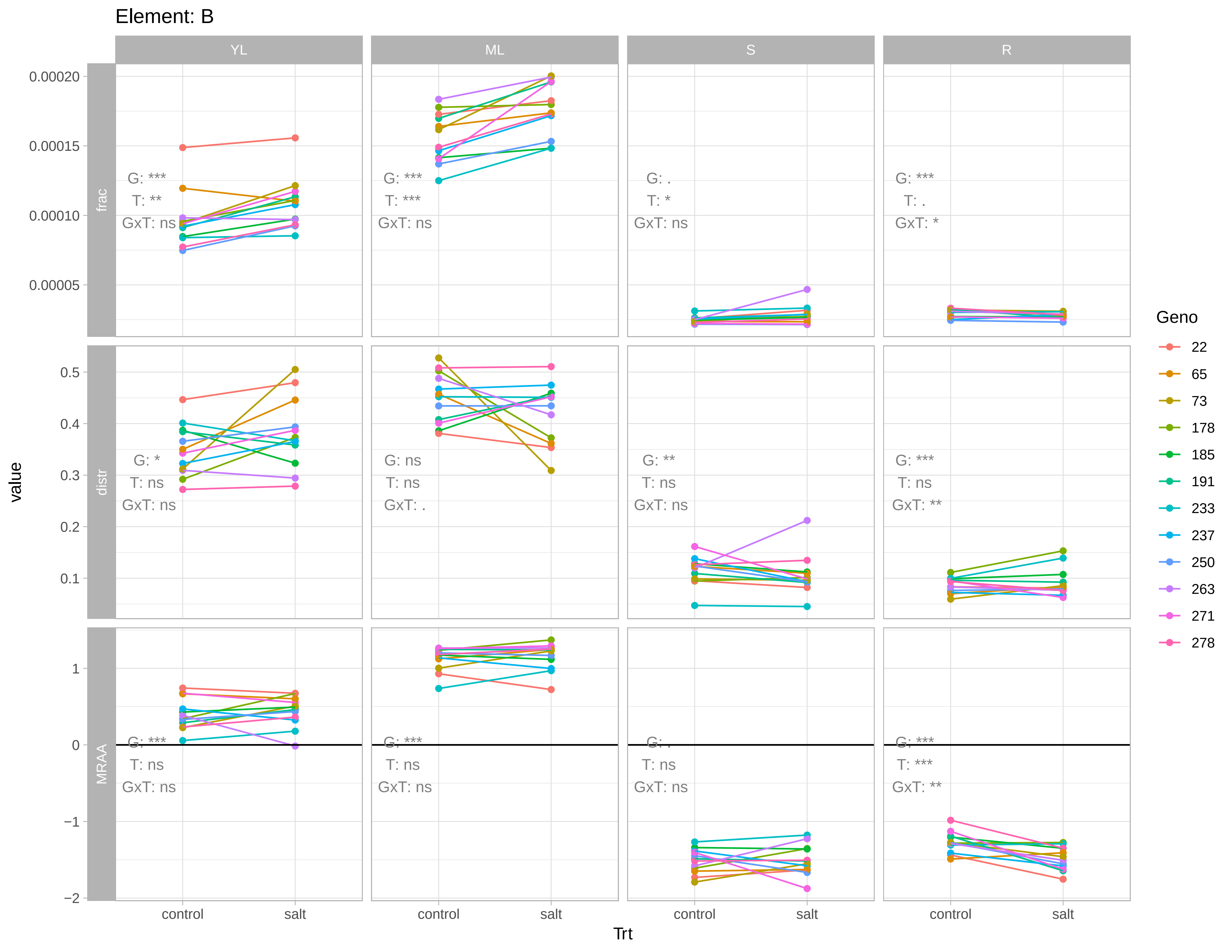

Element: B
YL
ML
S
R
frac
distr
MRAA
0.00020
0.00015
0.00010
0.00005
0.5
0.4
0.3
0.2
0.1
1
0
−1
−2
G: ***
T: **
GxT: ns
G: ***
T: ***
GxT: ns
G: .
T: *
GxT: ns
G: ***
T: .
GxT: *
G: *
T: ns
GxT: ns
G: ns
T: ns
GxT: .
G: **
T: ns
GxT: ns
G: ***
T: ns
GxT: **
G: ***
T: ns
GxT: ns
G: ***
T: ns
GxT: ns
G: .
T: ns
GxT: ns
G: ***
T: ***
GxT: **
value
control
salt
Trt
Geno
22
65
73
178
185
191
233
237
250
263
271
278

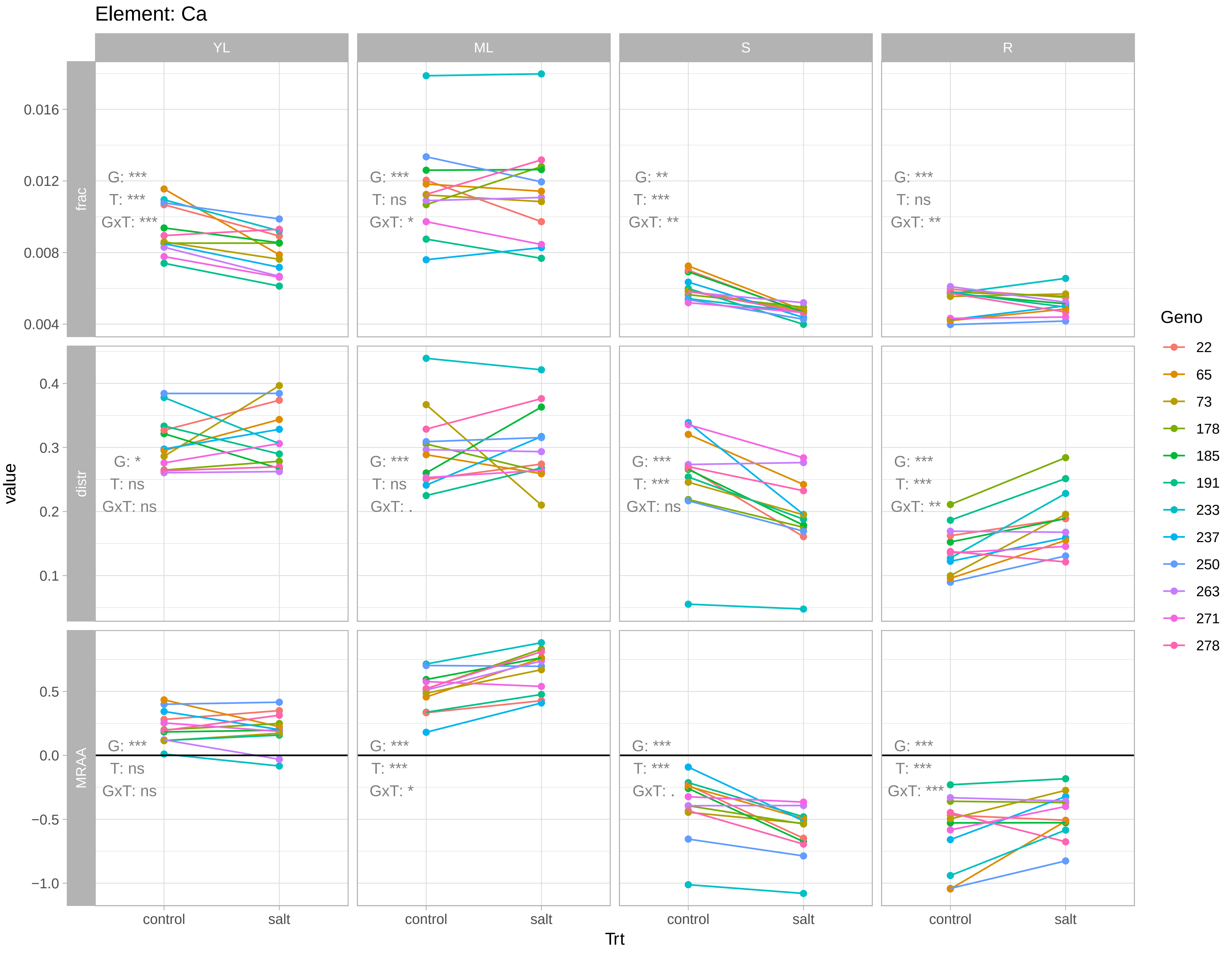

Element: Ca
YL
ML
S
R
frac
distr
MRAA
G: ***
T: ***
GxT: ***
G: ***
T: ns
GxT: *
G: **
T: ***
GxT: **
G: ***
T: ns
GxT: **
G: *
T: ns
GxT: ns
G: ***
T: ns
GxT: .
G: ***
T: ***
GxT: ns
G: ***
T: ***
GxT: **
G: ***
T: ns
GxT: ns
G: ***
T: ***
GxT: *
G: ***
T: ***
GxT: .
G: ***
T: ***
GxT: ***
0.004
0.008
0.012
0.016
0.1
0.2
0.3
0.4
−1.0
−0.5
0.0
0.5
value
control
salt
Trt
Geno
22
65
73
178
185
191
233
237
250
263
271
278

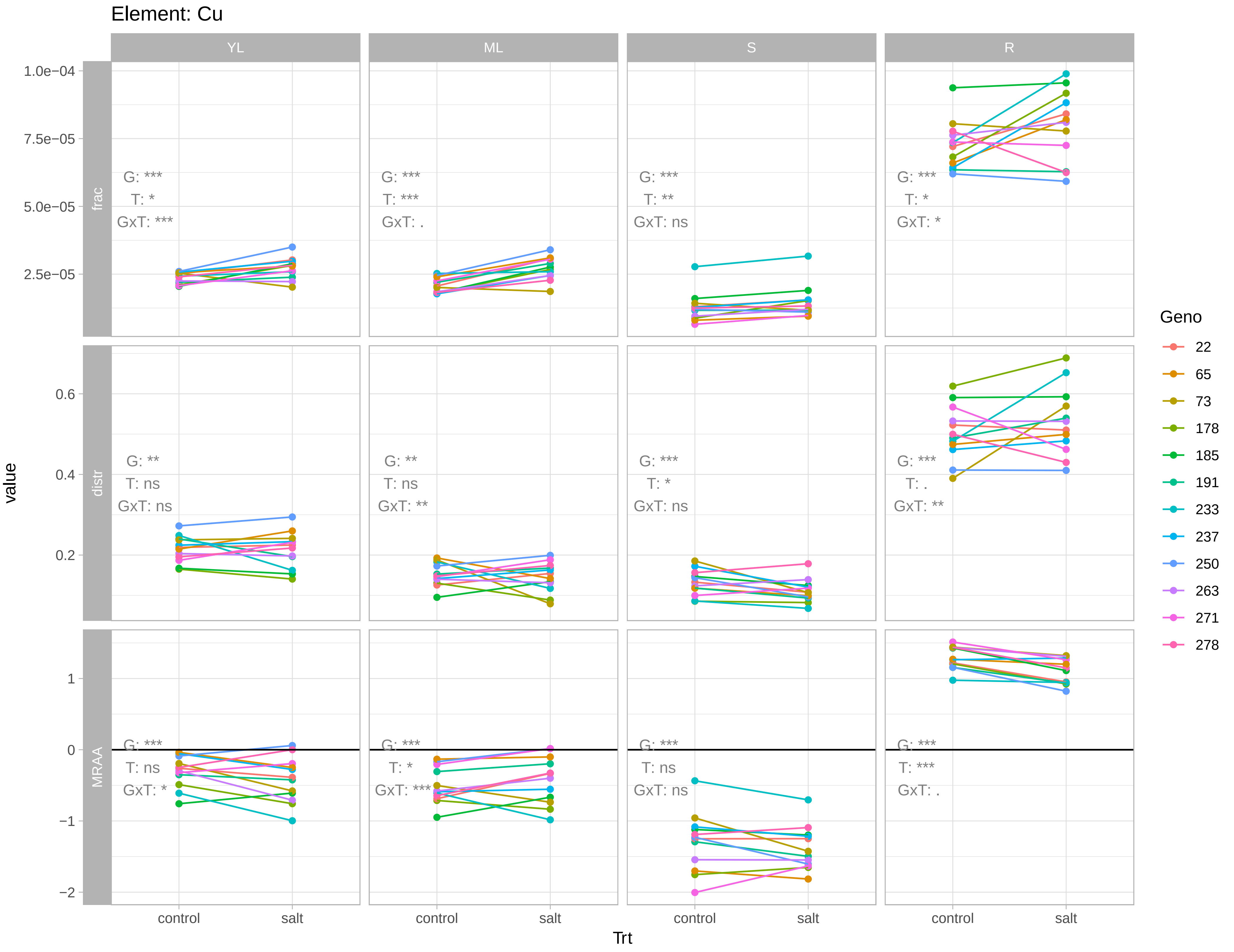
Element: Cu
YL
ML
S
R
frac
distr
MRAA
value
Trt
control
salt
G: ***
T: *
GxT: ***
G: ***
T: ***
GxT: .
G: ***
T: **
GxT: ns
G: ***
T: *
GxT: *
G: **
T: ns
GxT: ns
G: **
T: ns
GxT: **
G: ***
T: *
GxT: ns
G: ***
T: .
GxT: **
G: ***
T: ns
GxT: *
G: ***
T: *
GxT: ***
G: ***
T: ns
GxT: ns
G: ***
T: ***
GxT: .
Geno
22
65
73
178
185
191
233
237
250
263
271
278

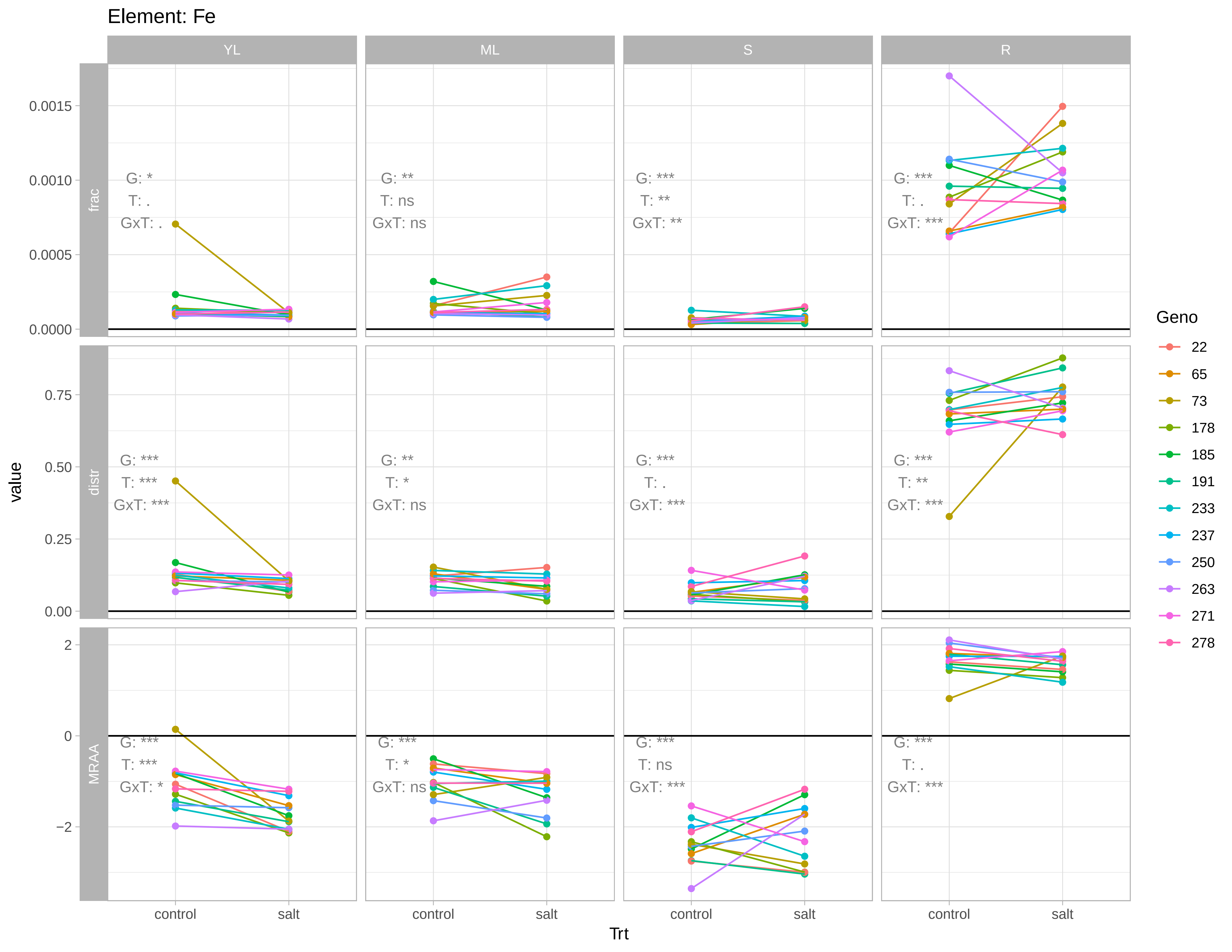

Element: Fe
YL
ML
S
R
frac
distr
MRAA
value
Trt
control
salt
G: *
T: .
GxT: .
G: **
T: ns
GxT: ns
G: ***
T: **
GxT: **
G: ***
T: .
GxT: ***
G: ***
T: ***
GxT: ***
G: **
T: *
GxT: ns
G: ***
T: .
GxT: ***
G: ***
T: **
GxT: ***
G: ***
T: ***
GxT: *
G: ***
T: *
GxT: ns
G: ***
T: ns
GxT: ***
G: ***
T: .
GxT: ***
Geno
22
65
73
178
185
191
233
237
250
263
271
278

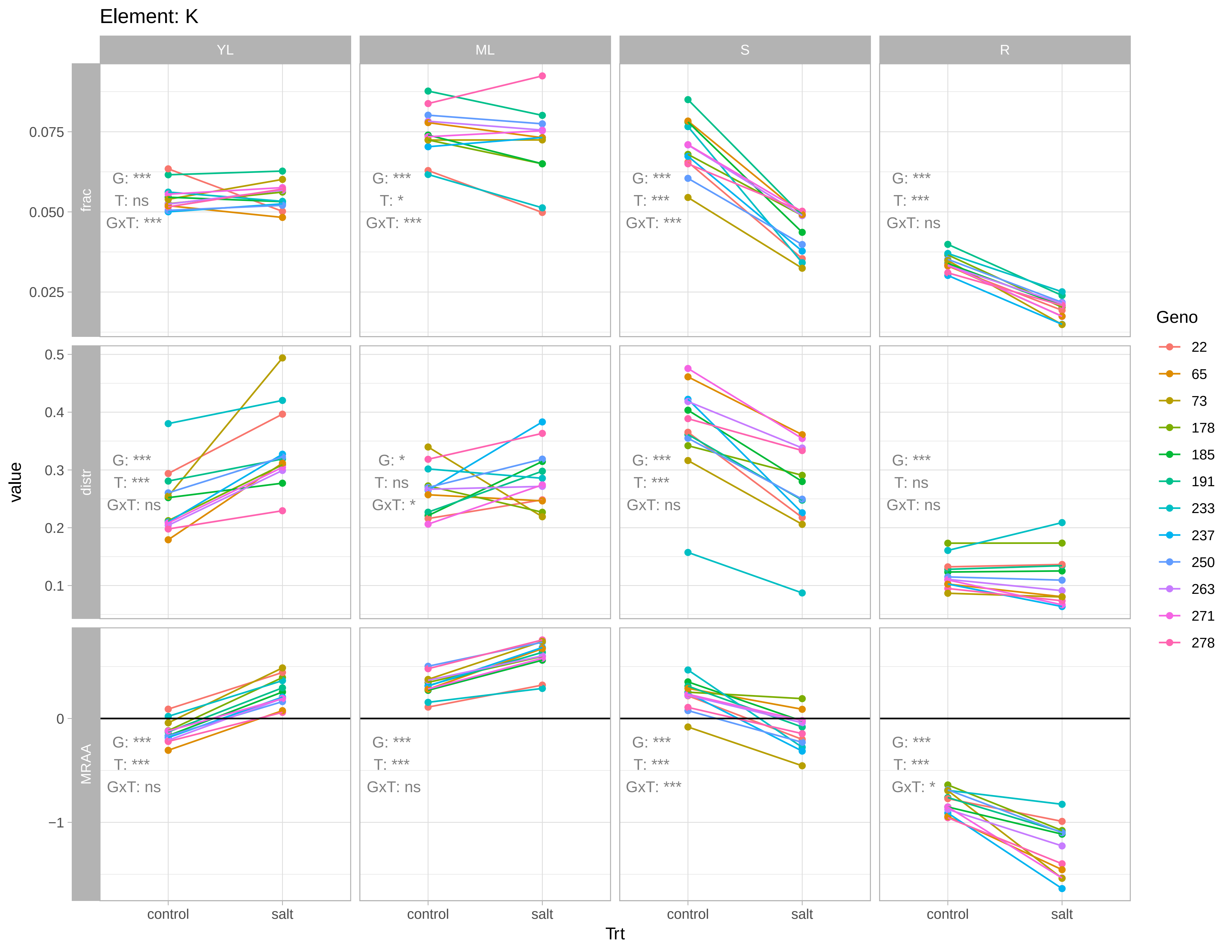
Element: K
YL
ML
S
R
frac
distr
MRAA
value
Trt
control
salt
0.075
0.050
0.025
0.5
0.4
0.3
0.2
0.1
0
−1
G: ***
T: ns
GxT: ***
G: ***
T: *
GxT: ***
G: ***
T: ***
GxT: ***
G: ***
T: ***
GxT: ns
G: ***
T: ***
GxT: ns
G: *
T: ns
GxT: *
G: ***
T: ***
GxT: ns
G: ***
T: ns
GxT: ns
G: ***
T: ***
GxT: ns
G: ***
T: ***
GxT: ns
G: ***
T: ***
GxT: ***
G: ***
T: ***
GxT: *
Geno
22
65
73
178
185
191
233
237
250
263
271
278

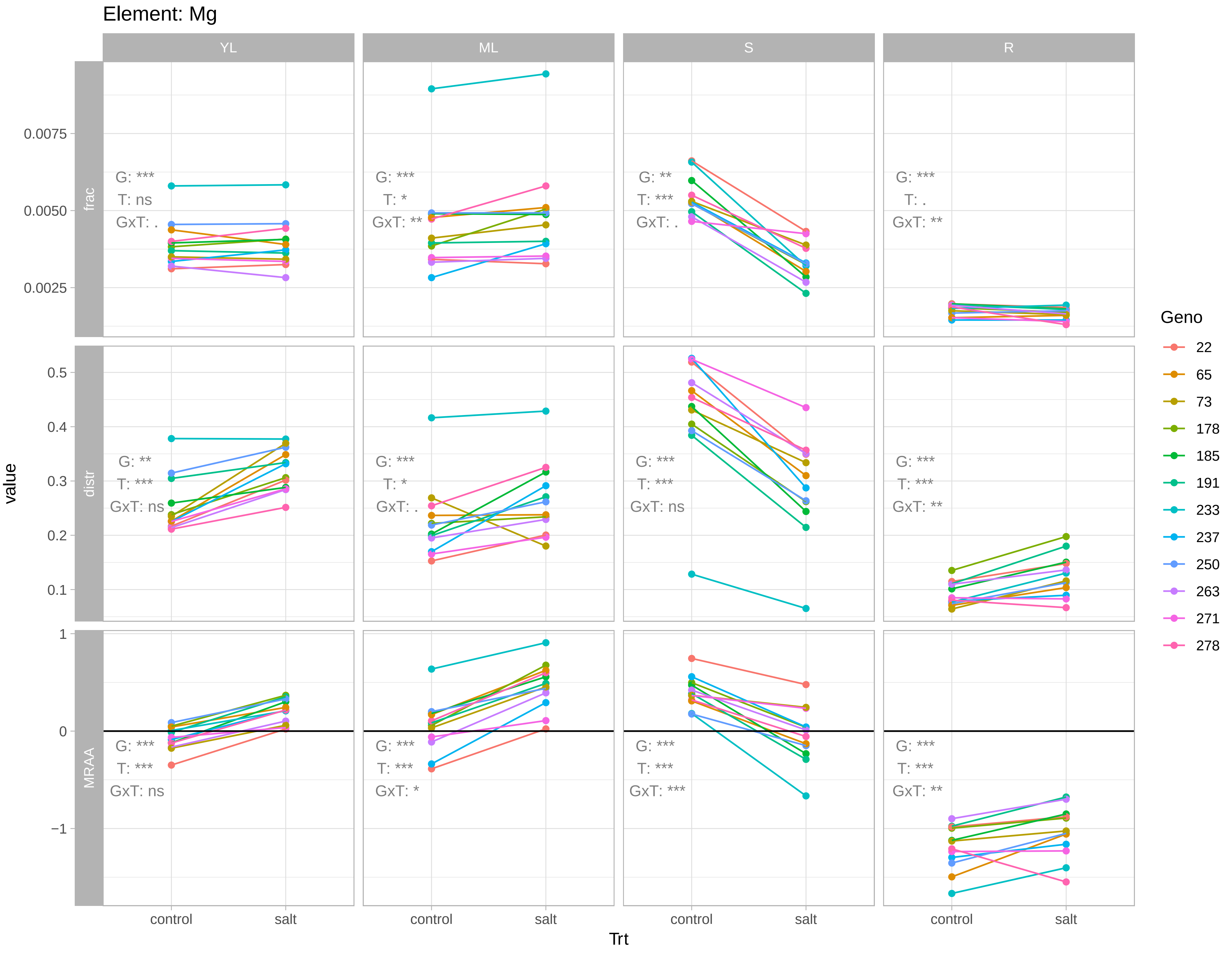

Element: Mg
YL
ML
S
R
frac
distr
MRAA
value
Trt
control
salt
0.0075
0.0050
0.0025
0.5
0.4
0.3
0.2
0.1
1
0
−1
G: ***
T: ns
GxT: .
G: ***
T: *
GxT: **
G: **
T: ***
GxT: .
G: ***
T: .
GxT: **
G: **
T: ***
GxT: ns
G: ***
T: *
GxT: .
G: ***
T: ***
GxT: ns
G: ***
T: ***
GxT: **
G: ***
T: ***
GxT: ns
G: ***
T: ***
GxT: *
G: ***
T: ***
GxT: ***
G: ***
T: ***
GxT: **
Geno
22
65
73
178
185
191
233
237
250
263
271
278

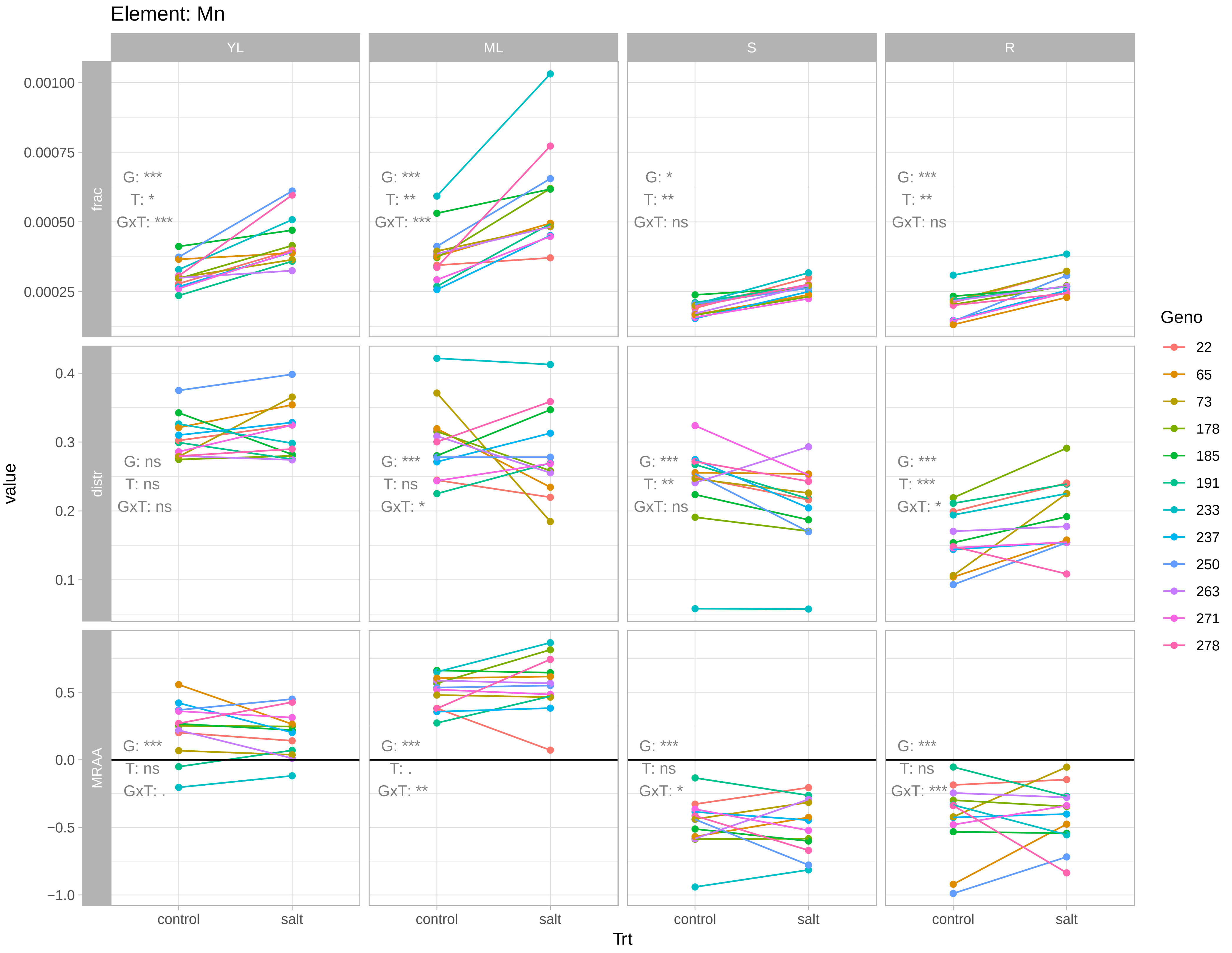

Element: Mn
YL
ML
S
R
frac
distr
MRAA
value
Trt
control
salt
Geno
22
65
73
178
185
191
233
237
250
263
271
278
0.00100
0.00075
0.00050
0.00025
0.4
0.3
0.2
0.1
0.5
0.0
−0.5
−1.0
G: ***
T: *
GxT: ***
G: ***
T: **
GxT: ***
G: *
T: **
GxT: ns
G: ***
T: **
GxT: ns
G: ns
T: ns
GxT: ns
G: ***
T: ns
GxT: *
G: ***
T: **
GxT: ns
G: ***
T: ***
GxT: *
G: ***
T: ns
GxT: .
G: ***
T: .
GxT: **
G: ***
T: ns
GxT: *
G: ***
T: ns
GxT: ***

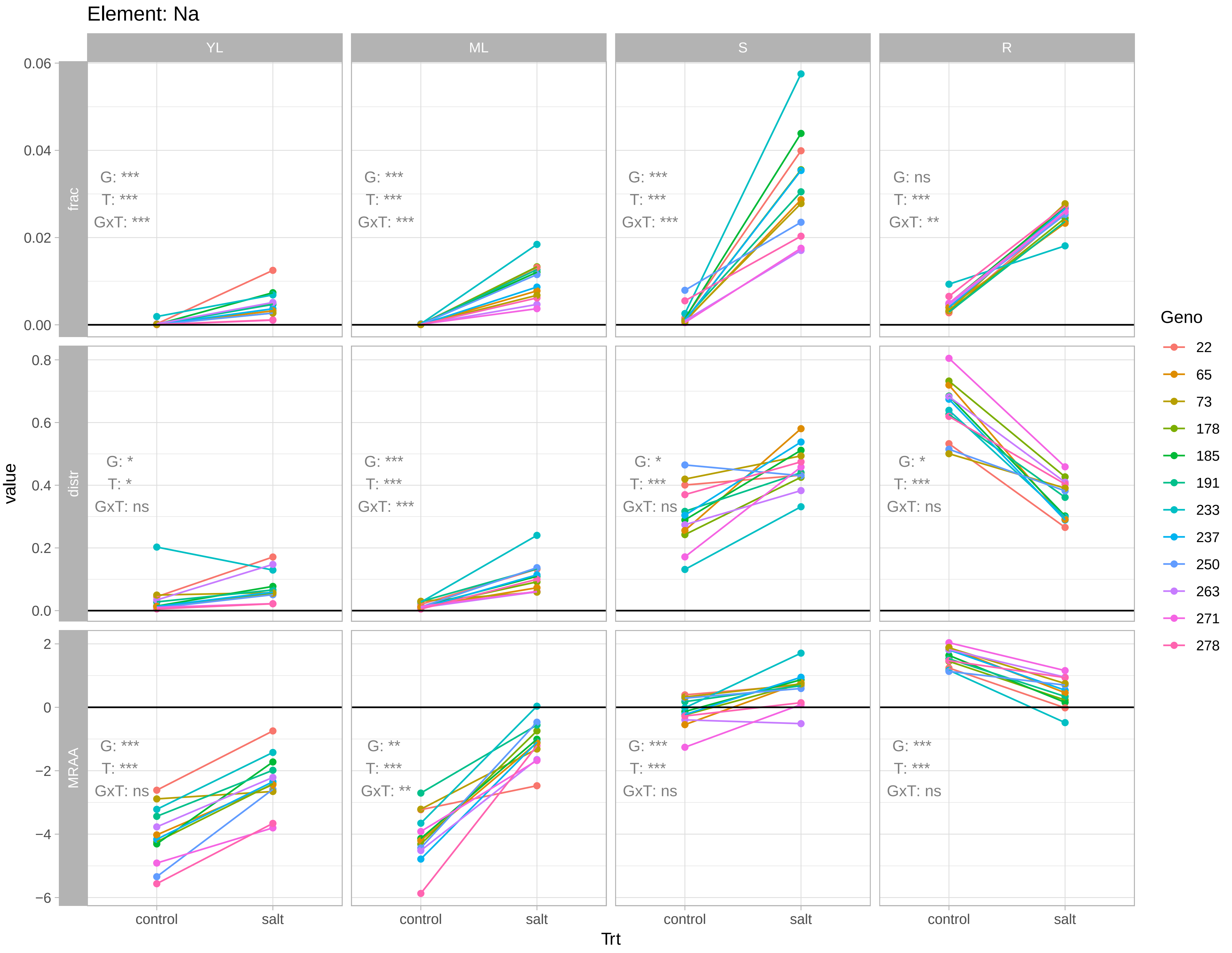

Element: Na
YL
ML
S
R
frac
distr
MRAA
value
Trt
control
salt
G: ***
T: ***
GxT: ***
G: ns
T: ***
GxT: **
G: *
T: *
GxT: ns
G: ***
T: ***
GxT: ***
G: *
T: ***
GxT: ns
G: *
T: ***
GxT: ns
G: ***
T: ***
GxT: ns
G: **
T: ***
GxT: **
G: ***
T: ***
GxT: ns
G: ***
T: ***
GxT: ns
Geno
22
65
73
178
185
191
233
237
250
263
271
278

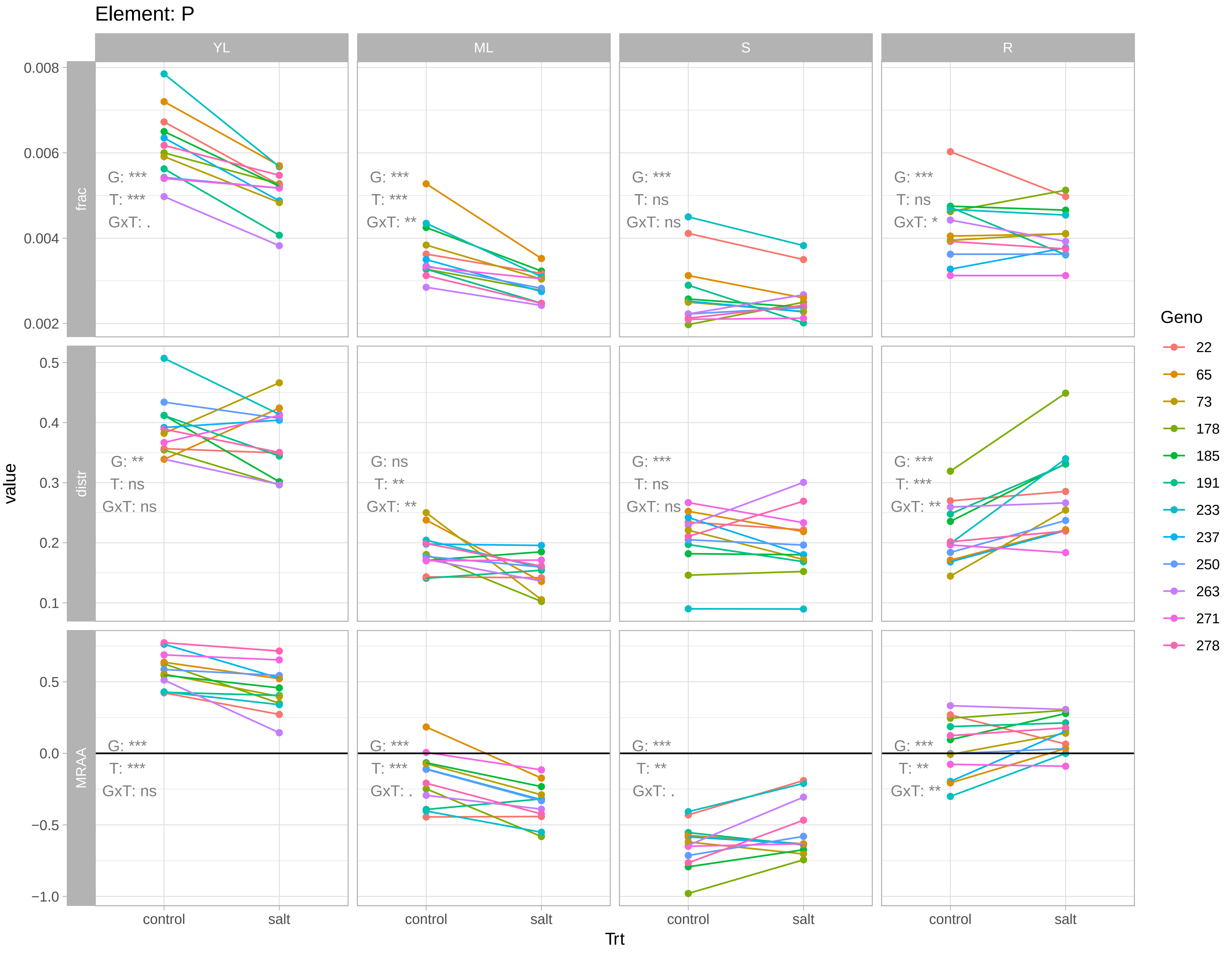

Element: P
YL
ML
S
R
frac
distr
MRAA
value
Trt
control
salt
Geno
22
65
73
178
185
191
233
237
250
263
271
278
G: ***
T: ***
GxT: .
G: ***
T: ***
GxT: **
G: ***
T: ns
GxT: ns
G: ***
T: ns
GxT: *
G: **
T: ns
GxT: ns
G: ns
T: **
GxT: **
G: ***
T: ns
GxT: ns
G: ***
T: ***
GxT: **
G: ***
T: ***
GxT: ns
G: ***
T: ***
GxT: .
G: ***
T: **
GxT: .
G: ***
T: **
GxT: **

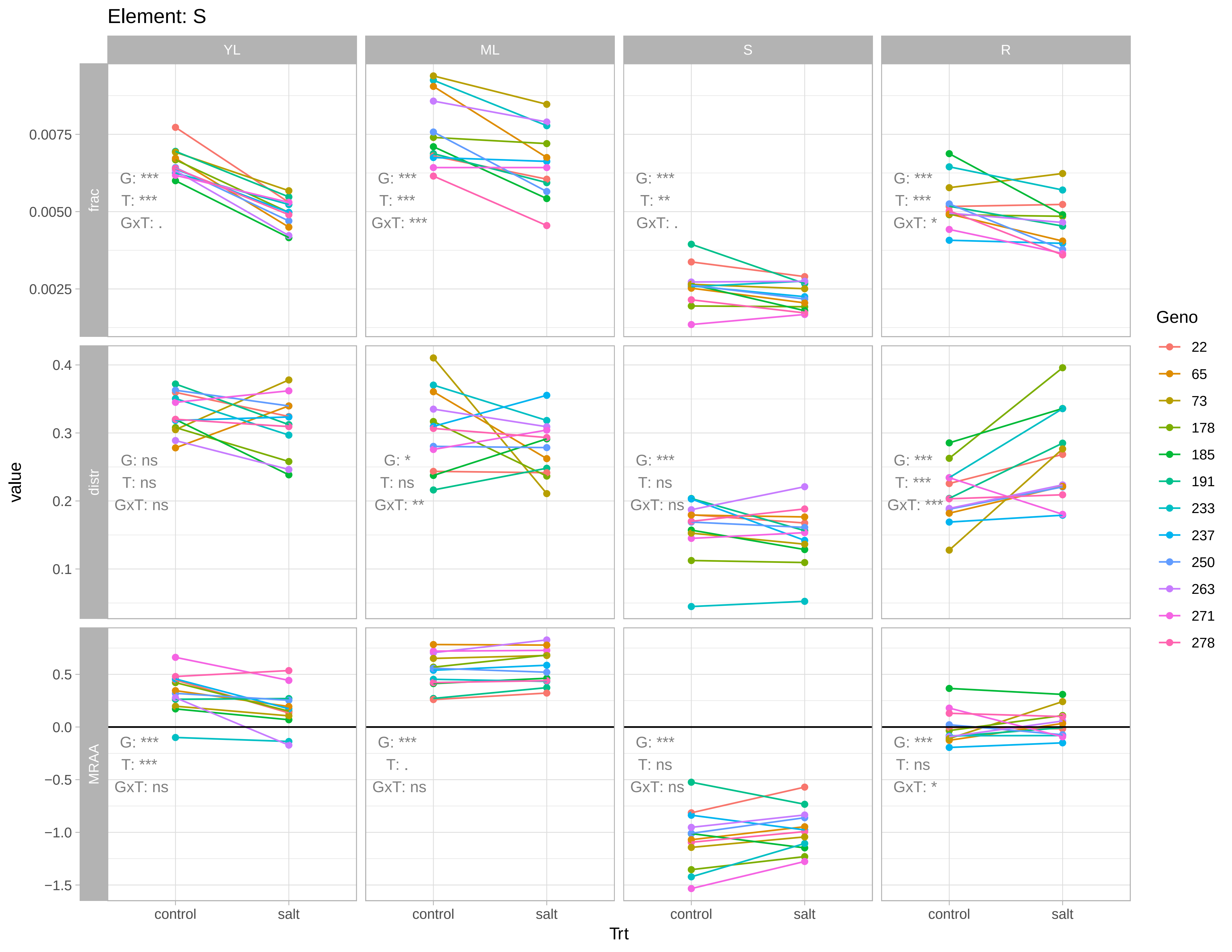
Element: S
YL
ML
S
R
frac
distr
MRAA
value
Trt
control
salt
Geno
22
65
73
178
185
191
233
237
250
263
271
278
G: ***
T: ***
GxT: .
G: ***
T: ***
GxT: ***
G: ***
T: **
GxT: .
G: ***
T: ***
GxT: *
G: ns
T: ns
GxT: ns
G: *
T: ns
GxT: **
G: ***
T: ns
GxT: ns
G: ***
T: ***
GxT: ***
G: ***
T: ***
GxT: ns
G: ***
T: .
GxT: ns
G: ***
T: ns
GxT: ns
G: ***
T: ns
GxT: *
0.0075
0.0050
0.0025
0.4
0.3
0.2
0.1
0.5
0.0
−0.5
−1.0
−1.5

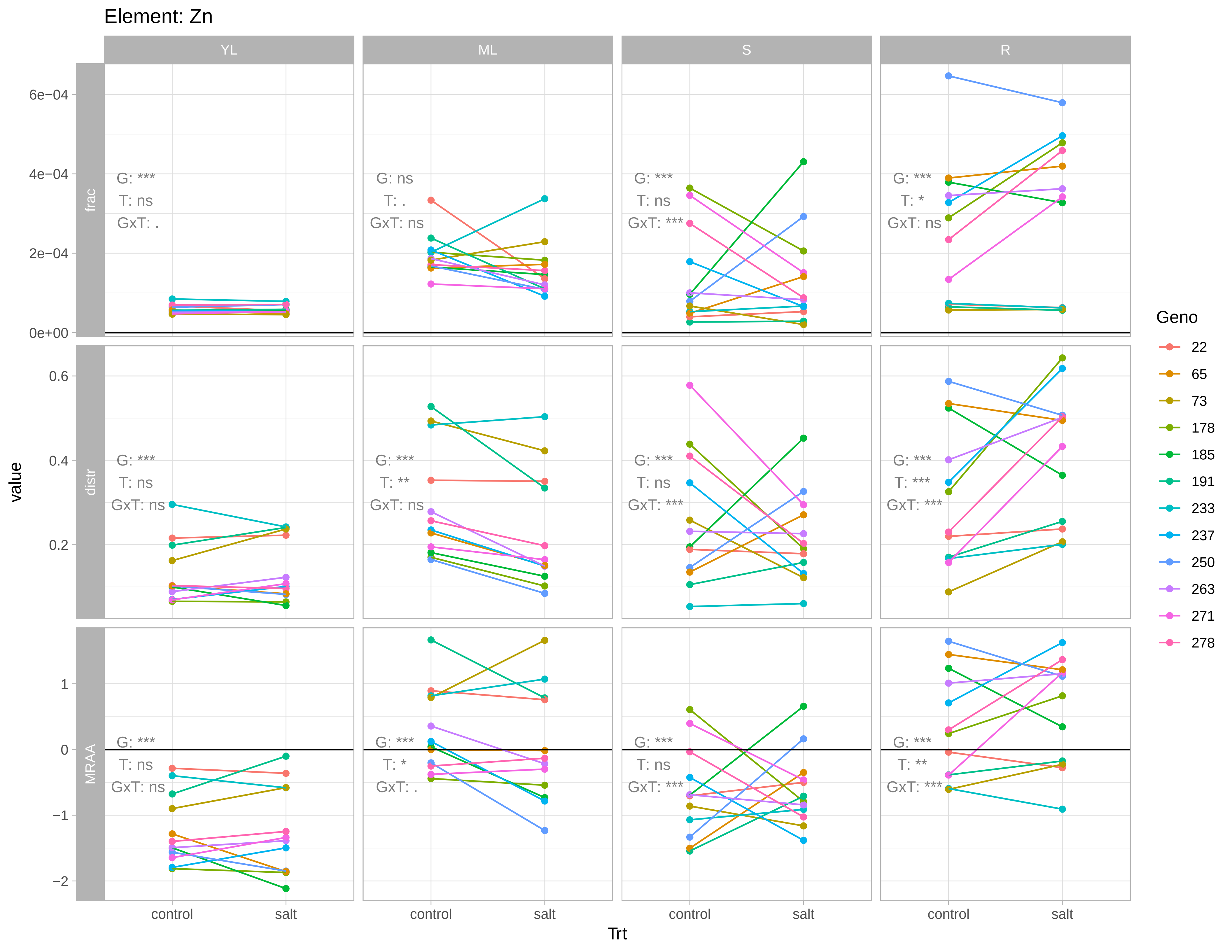

Element: Zn
YL
ML
S
R
frac
distr
MRAA
value
Trt
control
salt
6e−04
4e−04
2e−04
0e+00
0.6
0.4
0.2
1
0
−1
−2
G: ***
T: ns
GxT: .
G: ns
T: .
GxT: ns
G: ***
T: ns
GxT: ***
G: ***
T: *
GxT: ns
G: ***
T: ns
GxT: ns
G: ***
T: **
GxT: ns
G: ***
T: ns
GxT: ***
G: ***
T: ***
GxT: ***
G: ***
T: ns
GxT: ns
G: ***
T: *
GxT: .
G: ***
T: ns
GxT: ***
G: ***
T: **
GxT: ***
Geno
22
65
73
178
185
191
233
237
250
263
271
278
